# Using Multilayer Heterogeneous Networks to Infer Functions of Phosphorylated Sites

**DOI:** 10.1101/2020.08.25.266072

**Authors:** Joanne Watson, Jean-Marc Schwartz, Chiara Francavilla

## Abstract

1

Mass spectrometry-based quantitative phosphoproteomics has become an essential approach in the study of cellular processes such as signaling. Commonly used methods to analyze phosphoproteomics datasets depend on generic, gene-centric annotations such as Gene Ontology terms which do not account for the function of a protein in a particular phosphorylation state. Analysis of phosphoproteomics data is hampered by a lack of phosphorylated site-specific annotations. We propose a method that combines shotgun phosphoproteomics data, protein-protein interactions, and functional annotations into a heterogeneous multilayer network. Phosphorylation sites are associated to potential functions using a random walk on heterogeneous network (RWHN) algorithm. We validated our approach against a model of the MAPK/ERK pathway and functional annotations from PhosphoSite Plus and were able to associate differentially regulated sites on the same proteins to their previously described specific functions. We further tested the algorithm on three previously published datasets and were able to reproduce their experimentally validated conclusions and to associate phosphorylation sites with known functions based on their regulatory patterns. Our approach provides a refinement of commonly used analysis methods and accurately predicts context-specific functions for sites with similar phosphorylation profiles.

**For table of contents only:** We confirm that the eTOC figure contains original material drawn by the authors.

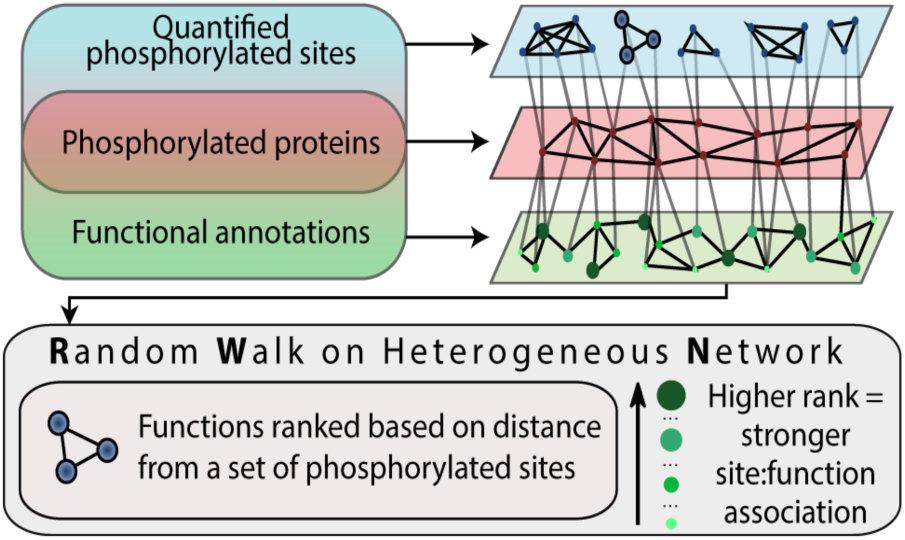

## 2. Introduction

Phosphorylation is the most studied post-translational modification (PTM) due to its central role in cellular regulation. It is thought to be the principal PTM in the human proteome, as well as an essential mediator of protein-protein interaction (PPI) and protein function ^1^. Transient changes occur at specifically regulated phosphorylation sites, of which there may be multiple on each protein. Regulation of phosphorylation is often dependent on perturbations such as the activity of extracellular ligands, drug treatment or physical stimuli in the extracellular environment ^2^. By comparing changes in the phosphoproteome of cells in different experimental conditions through mass spectrometry-based phosphoproteomics, phosphorylated sites that are key players in cellular processes and functions can be uncovered in an unbiased, high-throughput manner ^3^.

Functional analysis of phosphoproteomics datasets is typically based on gene-centric enrichment of Gene Ontology (GO) terms or involvement in known pathways ^4^. However, this approach disregards information captured by phosphoproteomics data on changes at specific phosphorylated sites, by limiting the analysis to the protein level. The modification state of a protein is inherently coupled to its function; PTMs alter protein activity, as well as the ability to interact with different sets of proteins. Furthermore, if a protein is phosphorylated on multiple sites, each with a different function and regulatory pattern, this information is not revealed by gene-centric analysis ^5^. For instance, the well-studied signalling protein MAPK1 has 18 known phosphorylated sites recorded in the database PhosphositePlus® (PSP), however only 6 have been annotated to a downstream function^6^. In the most recent release of PSP (v6.5.9.3), only 4271 of the more than 230,000 human phosphorylated sites recorded in the database are associated to 19 generic biological functions (e.g., “cell cycle”, “transcription”), which are qualified with one of: “induced”, “inhibited”, “regulated” or “altered”. Enrichment analyses that rely on generalizations based on protein-level or gene-centric descriptions exclude the details that are encoded in the phosphorylation signature. Analyses are thus hampered by the lack of phosphorylation site-specific functional annotations.

Several network-based methods have been proposed to move towards phosphorylation site-specific analyses of phosphoproteomics data. A significant focus has been on the inference of kinase-substrate networks, such as NetworKIN ^7^, KSEA ^8^, and IKAP ^9^, among others. These methods are useful for reconstructing the architecture of signalling intracellular networks, which can be informative for identifying modules of modified proteins involved in cellular processes, but again remain hampered by the lack of site-specific functional annotation ^10^. They may also be biased towards the most studied kinases and the exclusion of non-kinase proteins. Rudolph *et al.* ^11^ proposed a method to address this issue named PHOTON which identified differentially regulated proteins based on the level of phosphorylation of their binding partners in a high-confidence PPI network, and then used logistic regression to identify the involvement of the phosphorylated proteins in known signalling pathways. PHOTON is not truly site-centric, however, as it relies on summarized quantitative values of phosphorylated sites. An alternative approach described by Krug *et al.* ^12^ uses a modification to the gene-centric method gene set enrichment analysis (GSEA), which relies on a curated resource containing literature-derived phosphorylated site-specific signatures to assign functions to sites. However, the use of such a resource is limited by the lack of phosphorylated-site annotation to these signatures or specific cellular processes; though useful for identifying well-studied sites, predictions of functions of under-studied or sites in alternative contexts would not be captured. These do not fulfil the same role as popular gene-centric methods such as over-representation analysis (ORA)^13^ in aiding prediction and hypothesis generation when performing exploratory analysis of shotgun phosphoproteomics data.

In recent years, heterogeneous or multilayer networks have been used to represent many types of ‘omics datasets ^14^. These specialized networks are used to describe multiple types of associations with nodes representing different entities. To identify relationships between the different biological layers, random walk algorithms have been applied to these networks. The random walk on heterogeneous network (RWHN) with restart (RWR) method ^15^ has been particularly popular. Jiang ^16^ used RWR to prioritize disease candidate genes in a PPI-phenome network; similarly, Soul *et al.* ^17^ applied it to a PPI-phenome network to identify disease mechanisms. Recent work has extended the method to multiple layers of biological information, for example to infer disease associated m6A RNA methylation via known gene-disease associations ^18^. Similar methodology was used to associate phosphorylation sites recorded in the PSP database to diseases via kinase-substrate interactions ^19^.

Here, we propose an algorithm that uses RWHN to associate phosphorylated sites to context-specific function via a heterogeneous multilayer network using shotgun phosphoproteomics data. The network combines three layers of information: phosphorylated sites, protein interactions and GO terms. Clustering of phosphoproteomics data is used to find common features within datasets and is generally followed by enrichment analyses. This is based on the assumption that common patterns of phosphorylation, based on temporal changes or in response to a particular stimulus, treatment or environmental context, are a likely indicator of involvement in common functions or processes ^20^. We utilize this concept in our algorithm by connecting phosphorylated sites that have been clustered together, and therefore share regulatory patterns, within the multilayer network. The algorithm is intended for use on large phosphoproteomics datasets, assessing perturbations or processes as opposed to an interpretation of the full phosphoproteomics network in the cell.

We first apply our algorithm to a small-scale, manually curated validation network. We also assess the ability of our method and the most commonly used alternative, ORA, to capture the functional descriptions recorded in PSP, which are based on experimental analysis. We then demonstrate the utility of our algorithm using three previously published shotgun phosphoproteomics datasets, describing early signalling events in HeLa cells upon EGF and TGF-α stimulation ^21^, phosphorylation-mediated changes in breast cancer cells resistant to lapatinib treatment ^22^ and subcellular location dependent signalling events downstream of HRAS ^23^, respectively. We demonstrate that phosphorylated sites can be differentially assigned to functional annotations and this is driven by changes in the context-dependent modification of these sites. Our method is suitable for use with any phosphoproteomics dataset and could be generalized for data describing other PTMs.

## 3. Experimental Procedures

A multilayer heterogeneous network was constructed to relate biological function to phosphorylation sites via a protein-protein interaction network (Figure 1). Three types of nodes are contained within the network: phosphorylation sites (called ‘sites’ here on in for brevity), proteins and functional annotations (either Gene Ontology Biological Process (GOBP) terms or KEGG pathways). The edges describe five possible associations which are either bipartite (i.e., site to protein, protein to function) or between the same type of node.

**Figure 1.**
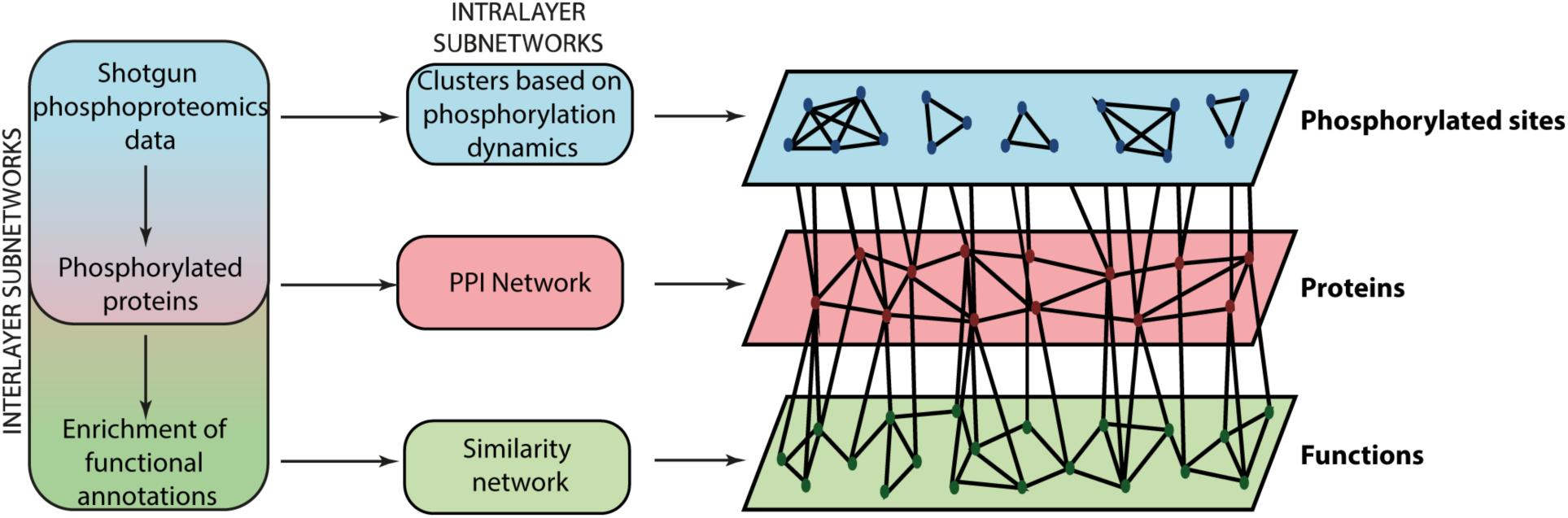
Overview of multilayer heterogeneous network construction from phosphoproteomics data.

### Dataset Accession and Processing

Sites annotated to have a role in biological processes were extracted from the ‘Regulatory sites’ dataset, available for download from PhosphoSitePlus® (version 6.5.9.3). The dataset was filtered on the “ON_PROCESS” column to only include sites annotated to biological roles, rather than molecular or interaction roles.

The results of the phosphoproteomics experiment described in Francavilla *et al.* ^21^ were taken from Supplementary Table S1 of their manuscript. The authors had processed the raw mass spectrometry data using MaxQuant ^24^, filtered identified phosphorylated sites based on a localization probability of greater than 0.75 and normalized remaining data to the ratio of EGFR at 1 min after stimulation with EGF or TGF-α. Prior to multilayer heterogeneous network construction, the data were further filtered to remove sites with two or more missing ratios in their time series and containing at least one SILAC ratio higher than 2 or lower than 0.5 (as described in the methods of Francavilla *et al.* ^21^). Missing data were imputed with random draws from a truncated distribution, as previously described ^25^, using the *impute.QRLIC* function from the *imputeLCMD* R package.

Data from the Ruprecht *et al.* ^22^ study were taken from the Supplementary Data file named ‘Filtered and normalized phosphoproteome dataset’. The raw mass-spectrometry data had been processed using MaxQuant, filtered to include only identified phosphorylated sites with a localization probability of greater than 0.75, and normalized. Sites that did not show a significant change (FDR <1%) either between the untreated parental and lapatinib-treated resistant (SILAC ratios H/L) or between the lapatinib treated parental and lapatinib-treated resistant conditions (SILAC ratios H/M) experimental conditions were filtered from the data, as described in the methods of Ruprecht *et al.* ^22^. Missing data were imputed with the same method used for the Francavilla *et al*. dataset.

Data from the Santra *et al*. study ^23^ were taken from Supplemental table S2. The raw mass-spectrometry data had been processed using MaxQuant, filtered to include only identified phosphorylated sites with a localization probability of greater than 0.75; significant sites were identified using a two-sample t-test with FDR. The downloaded dataset was normalized using the *limma* R package function *normalizeBetweenArrays* using the quantile method ^26^. Missing data were imputed with the same method used for the Francavilla *et al*. dataset. The data were then filtered to exclude sites that had not been identified by significant in the original analysis by Santra *et al*. Before applying the algorithm, all datasets were clustered based on what was reported in the original publication, where available. The model dataset was clustered using the Fuzzy C-Means (FCM) method ^27^, with the number of clusters selected based on the temporal trends we could visually identify in the data; generally, silhouette analysis is recommended for selecting the number of clusters using FCM for real experimental datasets ^28^. The sites extracted from PSP were clustered based on what process they were annotated to (as indicated in the “ON_PROCESS” column in the Regulatory_sites file available for download on the PSP website). The regulated phosphorylated sites from Francavilla et al. were clustered using FCM (Figure 3); since the original publication used a cluster number of 6, the same was done here for better comparison. The regulated phosphorylated sites from Ruprecht et al. and Santra et al. were both clustered using the k-means method, with the cluster number selected using the elbow method (Figure S4 and S5, respectively) ^29^.

### Construction of Multilayer Network

#### Phosphorylation Site – Phosphorylation Site Subnetwork

Edges were drawn between sites in the same cluster that had a Pearson correlation (R2) between all the data points greater than or equal to 0.99 (Figure 1).

#### Protein - Protein Subnetwork

The protein-protein interaction network was extracted from STRING ^30^. All interactors of the proteins included in the phosphoproteomics datasets with an experimental confidence score of greater than 0.4 were included (Figure 1).

#### Function – Function Subnetwork

We used either GOBP terms or KEGG pathways as functional annotators in this work. If GOBP terms were included in the multilayer network, the *GOSemSim* package from Bioconductor ^31^ was used to calculate the semantic similarity of enriched GOBP terms. An edge was drawn between terms with semantic similarity greater than 0.7, as calculated using the Wang method included in the *goSim* function of *GOSemSim* ^31, 32^. In the case of KEGG pathways, edges were drawn between pathways that had greater than 70% pairwise similarity in their functional annotation profiles, following the method described in Stoney *et al.* ^33^ (Figure 1).

#### Phosphorylation Site – Protein Bipartite Edges

Sites and proteins had an edge between them if the residue was found on that protein. Therefore, sites will only have one edge, but proteins will have edges to all the sites found on that protein that were differentially regulated in the data set (Figure 1).

#### Protein – Function Bipartite Edge

We assumed that closely connected nodes in the protein-protein subnetwork would more likely be involved in similar biological processes. Therefore, we computed modules of the protein-protein subnetwork using the Louvain module-detection method ^34^. Proteins from each module were analyzed for enrichment (hypergeometric test with Benjamini-Hochberg correction, FDR < 0.05%) of functional annotations (from either the “GO_Biological_Process_2018” or “KEGG_2019_Human” libraries, included in enrichR and listed on the EnrichR website) using the *enrichR* ^35^ R interface. This increased the specificity of terms to be included in the network. When GO terms were included, high frequency (annotated to more than 5% of genes) and semantically redundant terms (similarity > 0.9) were filtered using the Bioconductor *GOSemSim* ^31^ and *GO*.*db* packages ^36^ (Figure 1).

### Random Walk on Heterogeneous Network

The heterogeneous network can be represented as an adjacency matrix as follows:

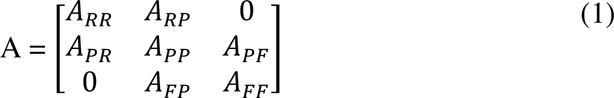

where: A_RR_ are site-site associations, A_PP_ are protein-protein associations, A_FF_ are function-function associations, A_RP_ are site-protein associations with A_PR_ as the transpose and A_PF_ are protein-function associations with A_FP_ as the transpose.

As described in previous work ^15^, a transition matrix (M) was calculated for use in the first stage of the algorithm:

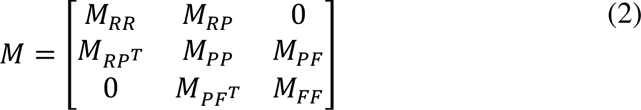

The bipartite inter-subgraph transition matrices (M_RP_ and M_PF_) were calculated as:

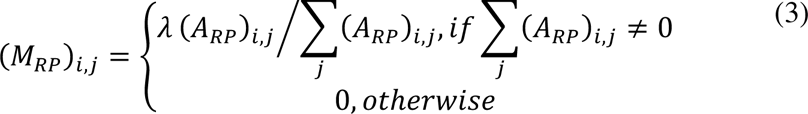

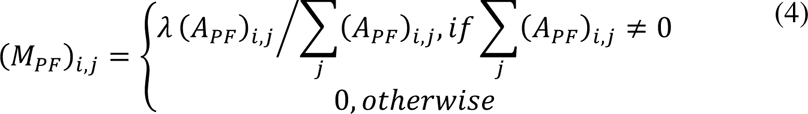

where λ is the transition probability (i.e., the likelihood of the walker moving between two layers of the network).

The intra-subgraph transition matrices (M_RR_, M_PP_ and M_FF_) were calculated as:

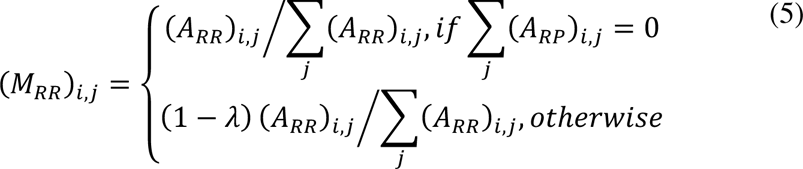

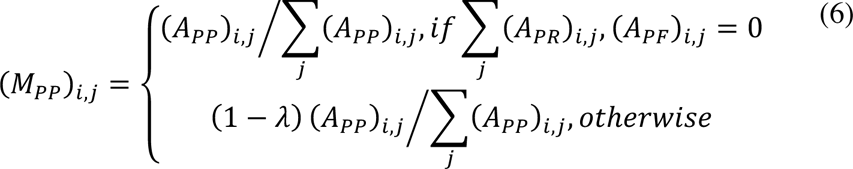

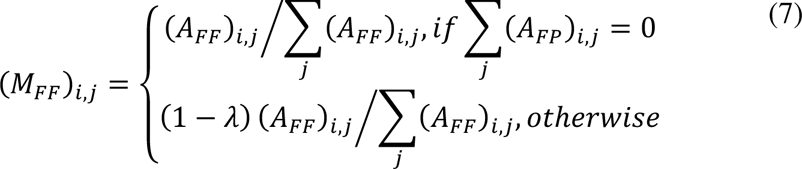

RWHN is a ranking algorithm; nodes are ranked based on the probabilities of finding the random walker at a given node in the steady state, having started at a given seed node or set of seed nodes. In this work we set the seed nodes to be those sites belonging to a particular cluster. The probability of finding the random walker at each node for each step is calculated based on the iterative equation:

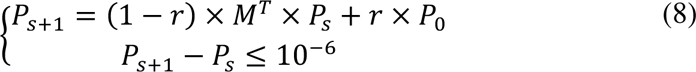

where *r* is the restart probability (set to 0.7 as described in Kohler *et al*. ^37^), P_0_ is the initial probability vector and P_s_ is the probability vector at step s. P0 was calculated such that all seed nodes were given equal probabilities with their sum equal to 1. All other nodes in the site-site subnetwork were assigned an initial probability of 0. Nodes in other subnetworks were assigned equal probabilities with their sum equal to 1 and weighted with the tunable parameters ηP and ηF, that are used to weight the influence of each layer.

The output of the algorithm is a ranked list of all the nodes, based on the probability of finding the random walker at each node in the steady state. This list is filtered by retaining functional annotations and removing proteins and sites. We remove annotations that are ranked in the same position regardless of seed node, in order to focus on specific functions related to each cluster. The top 5% of functions in the probability distribution are retained.

We implemented the algorithm in R using packages available from Bioconductor and CRAN. The source code is available at www.github.com/jowatson2011/RWHN_phosphoproteomics and an R package for general use of the tool can be found at github.com/JoWatson2011/phosphoR-WHN. RWHN on the validation network took less than 30 seconds to run however the larger experimental data sets took several hours on a moderately powerful computer (32GB RAM, Intel i7-4770 Processor).

### Overrepresentation Analysis (ORA)

The names of modified proteins in each cluster were used as input for the *enrichR* R interface ^35^. The libraries “GO_Biological_Process_2018” or “KEGG_2019_Human” (included in enrichR and listed on the EnrichR website) were used depending on which was used for the RWHN analysis of the same data. The list of overrepresented terms was filtered to only include those with an FDR < 0.05.

## 4. Results

### Overview of Algorithm

To associate phosphorylated sites of unknown function to potential cellular functions we developed an algorithm to apply to shotgun phosphoproteomics data. First, in order to help experimental biologists to identify the context- or perturbation-specific roles of modified proteins changing in each given experiment, a multilayer heterogeneous network is constructed based on the regulation of phosphorylated sites. Regulated phosphorylated sites may be defined differently by the user depending on the quantitation method used in their phosphoproteomics experiment (e.g., label-free quantification, SILAC) or their experimental question. The network represents three layers of biological entities and information: regulated phosphorylated sites, protein-protein interaction, and biological function (Figure 1). We then apply a ranking algorithm, RWHN, which ranks nodes of each layer based on: (i) distance from the phosphorylated sites of interest, which are assigned as ‘seed’ nodes and (ii) the topology of the multilayer heterogeneous network ^15^. Functions that are highly ranked can be considered more correlated with a set of seed nodes.

In the multilayer heterogeneous network, edges are drawn between sites based on similarity in the regulatory pattern of their phosphorylation which we determine using k-means or fuzzy C-means clustering. For the protein layer, a PPI network is constructed of the phosphorylated proteins and their interactors using the STRING database ^30^. Interactors are included to account for non-phosphorylated proteins, or those below the limit of detection of the experiment ^38^. To get a comprehensive and specific list of functional annotations, we identified closely connected groups of proteins in the PPI network using module detection and performed functional enrichment analysis on these modules. Edges between functional annotations are drawn based on functional similarity and overlap ^31, 33^.

The RWHN algorithm simulates a walker moving from a starting node(s) (called a seed) and then node to node through the multilayer network. Each step is influenced by the probability of transition to another layer (λ), the weighting of the protein and function layers (η_P_ and η_F_) and the probability of restart (that is, teleportation back to the seed node(s), *r*). The output of RWHN consists in a list of ranks for all the nodes in the network based on the likelihood of the walker reaching that node.

To optimize the algorithm, we ran RWHN with each of these parameters (λ, r, η_P_ and η_F_) tested over a range of values (0.1 – 0.9), changing one whilst setting all others to 0.5 (Figure S1A). Performance was decidedly stable over the range of η_P_ and η_F_ however altering λ and *r* resulted in differential ranking depending on the parameter value. Previous work has suggested an ideal λ and *r* of 0.7 ^15, 37, 39^, whilst η_P_ and η_F_ were set to 0.7 and 0.3 respectively to prioritize movement in the protein layer and reduce the number of terms having the same rank regardless of seed.

The parameters used in the construction of the PPI network were also assessed. Proteins interacting with the phosphorylated proteins found in a given sample were extracted from STRING; those proteins whose interaction had a STRING experimental confidence score above a given threshold were included in the multilayer heterogeneous network. To investigate the impact of choosing different STRING confidence scores on the results, we tested a confidence score range of 0.1 – 0.9. First, we looked at whether connectivity of the graph was altered by the score. As expected, the greater the confidence score, the more components the PPI were split into; above a score of 0.7, the PPI had two or more components (Figure S1B). We next investigated whether the RWHN results were affected by changing the STRING confidence score increasing components within the protein layer. Using weighted-tau correlation^40^ to compare the rankings from each set of results, we found that the correlation was generally in the range of 0.8 – 1.0, with the lowest value being 0.54 between networks constructed with 0.1 or 0.9 confidence threshold, which were the extremes of the range tested.

### Validation 1: Model of MAPK/ERK pathway

MAPK/ERK signalling has been well studied and the temporal phosphorylation status of component proteins in response to growth factor stimulus are relatively established ^41, 42^(Figure 2A). Based on this general understanding, we compiled a simple validation dataset of phosphorylation dynamics at particular sites of the main signalling proteins within the pathway (Figure 2B). Sites were chosen based on whether their phosphorylation is known to activate or inactivate protein activity, as recorded in the PSP database ^6^. We clustered the data into 5 clusters, referred to here as clusters 1-5, using the fuzzy c-means algorithm (Figure 2B). The features of the multilayer network that was constructed are summarized in Table 1. Enrichment of GOBP terms was used to form the functional annotation layer of the network.

**Figure 2.**
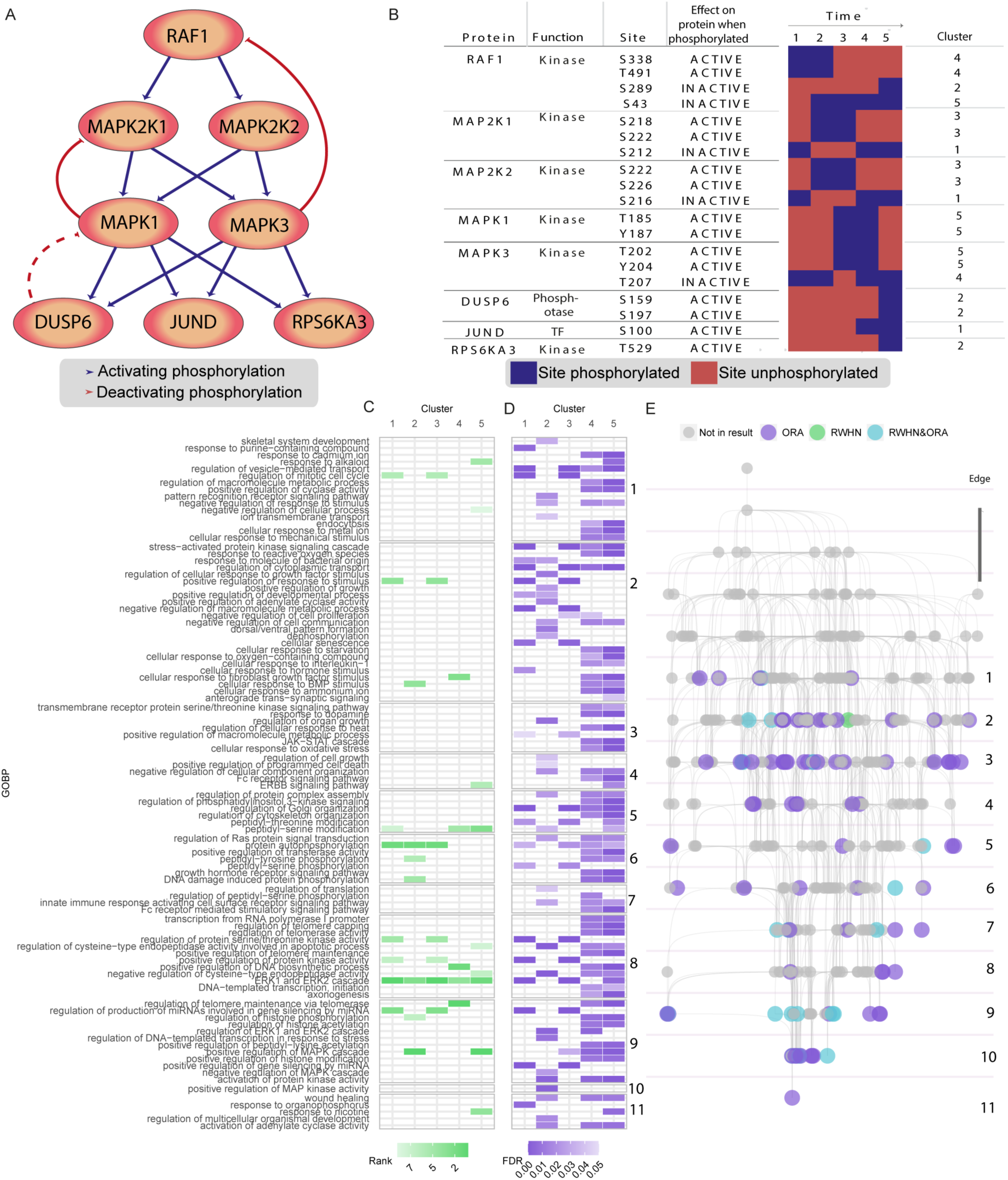
Site- and gene-centric analysis of validation data. A) Construction of validation data was based on a traditional representation of the MAPK/ERK pathway. B) A validation dataset was constructed based on the MAPK/ERK pathway, simulating phosphorylation dynamics over time after pathway activation. The sites were clustered using fuzzy c-means clusters. These data were used to construct a heterogeneous multilayer network. C) RWHN results with sites annotated to functional roles in PSP; seed nodes are equivalent to the clusters. D) ORA results with modified proteins contained in toy data. E) GO hierarchy, which has been subset to include only the parents of terms that occurred in either the RHWN or ORA analysis. Terms that occur in either set of results are colored in purple or green, respectively, with terms that occur in both colored in light blue.

**Table 1.** Multilayer heterogeneous network, constructed from MAPK/ERK validation data. GOBP terms were included as functional annotations.

| Subnetwork | Edges | Nodes |
| --- | --- | --- |
| Site-site | 28 | 19 |
| Protein-protein | 932 | 313 |
| Function-function | 148 | 88 |
| Site-protein | 19 | 19 sites, 8 proteins |
| Protein-function | 1796 | 261 proteins, 298 functions |

We ran the RWHN algorithm over the multilayer heterogeneous network with seed nodes set to all the sites belonging to one of the clusters; this was then repeated for each cluster (Figure 2C). The highest ranked GOBP term was the same (“protein autophosphorylation”) regardless of the seed nodes. However, it was possible to differentiate between the five clusters based on the ranking of terms below the first one.

Where available, we could use the biological process annotations in PSP (v6.5.9.3) as a ‘benchmark’ for the highly ranked terms associated to each cluster. RPS6KA3_Y529, annotated to “apoptosis, altered” in PSP is found in cluster 2, along with DUSP6 sites; this cluster highly ranks “positive regulation of programmed cell death”. Of the four RAF1 sites included in this model dataset, two were annotated to “cell cycle regulation” in PSP: RAF1_S289, in cluster 2, and RAF1_S338, in cluster 5. These two clusters both rank “regulation of mitotic cell cycle” in the top 5% of terms.

Largely due to the inherent step in our method in which the redundant terms are reduced in the multilayer heterogeneous network, fewer terms were found in the results of RWHN than ORA. We next wanted to assess whether the terms in the RWHN result were drawn from a particular level of the GO hierarchy; if they were, this would suggest a bias towards more specific or more broad terms compared to ORA. From the GO hierarchy, we extracted the parent and ancestor terms of the terms found in each set of the results (Figure 2E). We found that RWHN terms were drawn from the same range of the hierarchy as the ORA terms, with slightly more coming from lower in the hierarchy, indicating the results have more specificity than the ORA results. We note that there is also less redundancy in the RWHN results, despite the ORA results also being reduced for semantically similar / redundant terms. For example, the terms “regulation of ERK1 and ERK2 cascade” and “ERK1 and ERK2 cascade” both come up in the ORA results and are associated to different clusters.

To assess the robustness of the algorithm, we performed a random permutation control. RWHN was run one hundred times with the seeds set as before, random permutations of the subnetworks and bipartite edges maintained. Kernel Density Estimation (KDE) was calculated to assess how often each GO term occurred at each rank in the 100 random permutations for each set of seed nodes. In each case, there were a subset of terms that were more likely to have a rank greater than 50, however none strongly correlated with the actual rankings for each set of seed nodes (Figure S2A). This confirms that the rankings are not random, but primarily determined by the network topology. We next investigated the rate of potential false positives in the results. Previous implementations of RWHN, primarily using genomics data, have utilized an arbitrary threshold of the top *x* ranked terms, or the top *x*% of ranked terms (based on the probability vector as calculated in equation 8). A threshold that is too stringent would not remove lower-ranking terms from the results, whilst a too-low threshold would increase the amount of noise in the data. Using the results of RWHN run on the randomly permuted networks we calculated the probability that the “true” ranking (i.e. from RWHN run on the non-permuted network) was different from the random rankings (p < 0.05, Mann-Whitney U test with Benjamini-Hochberg adjustment ^43, 44^). We found that most of the rankings differed from random (Figure S2B). The terms that had p > 0.05 were checked against the RWHN results with different thresholds selected (top 1%, 5%, 10% and 15% of ranked terms); none were found above the 1% thresholds for any of the clusters, and only one term (“peptidyl-serine modification” in cluster 2) was found above the 5% and 10% thresholds (Figure S2B). We chose to set the default threshold to the top 5% of terms to be more conservative whilst retaining useful annotations.

For a given set of seed nodes, the algorithm is capable of giving high ranks to functions with known associations to those sites. It is also capable of predicting non-random, reasonable functions for sites of unknown function in this context.

### Validation 2: Classification of PhosphoSitePlus annotated sites

A comprehensive benchmark of dynamically regulated phosphorylated sites is not currently available, due to the lack of phosphorylated sites with a regulatory pattern universally defined independent of the experimental conditions. Therefore, we used the static classifications of phosphorylated site function recorded in PSP to benchmark the predictive power of our algorithm. The PSP database has annotations to biological processes for ∼4000 human phosphorylated sites. As these annotations are non-standard, each one was mapped to the most closely related GOBP term, where available (Table S1). The majority of sites were annotated to signal transduction, gene expression or cell differentiation (Figure 3A). Of these, 1496 sites were annotated to more than one function; as these annotations are manually assigned based on literature searches, the multiple functions may be due to the differences in experimental questions in the source datasets. The sites were clustered based on which function they were annotated to and analyzed with RWHN and ORA. The GOBP terms in the results of both methods were associated to the child terms of each mapped PSP GO term, per the GO hierarchy. The terms in the results of both methods showed a similar distribution to the terms in PSP as a whole, the main exception being in the cell differentiation category (Figure 3B). The two methods showed a considerable difference in the number of terms in the output, with ORA resulting in several hundred more terms in the output than RWHN. To test whether the two sets of results were drawn from the same parts of the GO hierarchy (with those from the top of the hierarchy being broader and those at the bottom being more specific), we constructed a tree based on the GO hierarchy. The tree was then divided into levels, as in Figure 2E, to capture terms that were of similar descriptiveness. Finally, we counted the number of terms that came up in the RWHN or ORA results (Figure 3C). Both sets of results draw from the same part of the hierarchy, with a skew towards the broader terms. We therefore concluded that RWHN resulted in as descriptive but more restricted and focused annotations than ORA. This was notable given that RWHN did not associate any terms to the smaller categories in PSP, such as exocytosis, endocytosis, cell growth and autophagy (Figure 3B).

**Figure 3.**
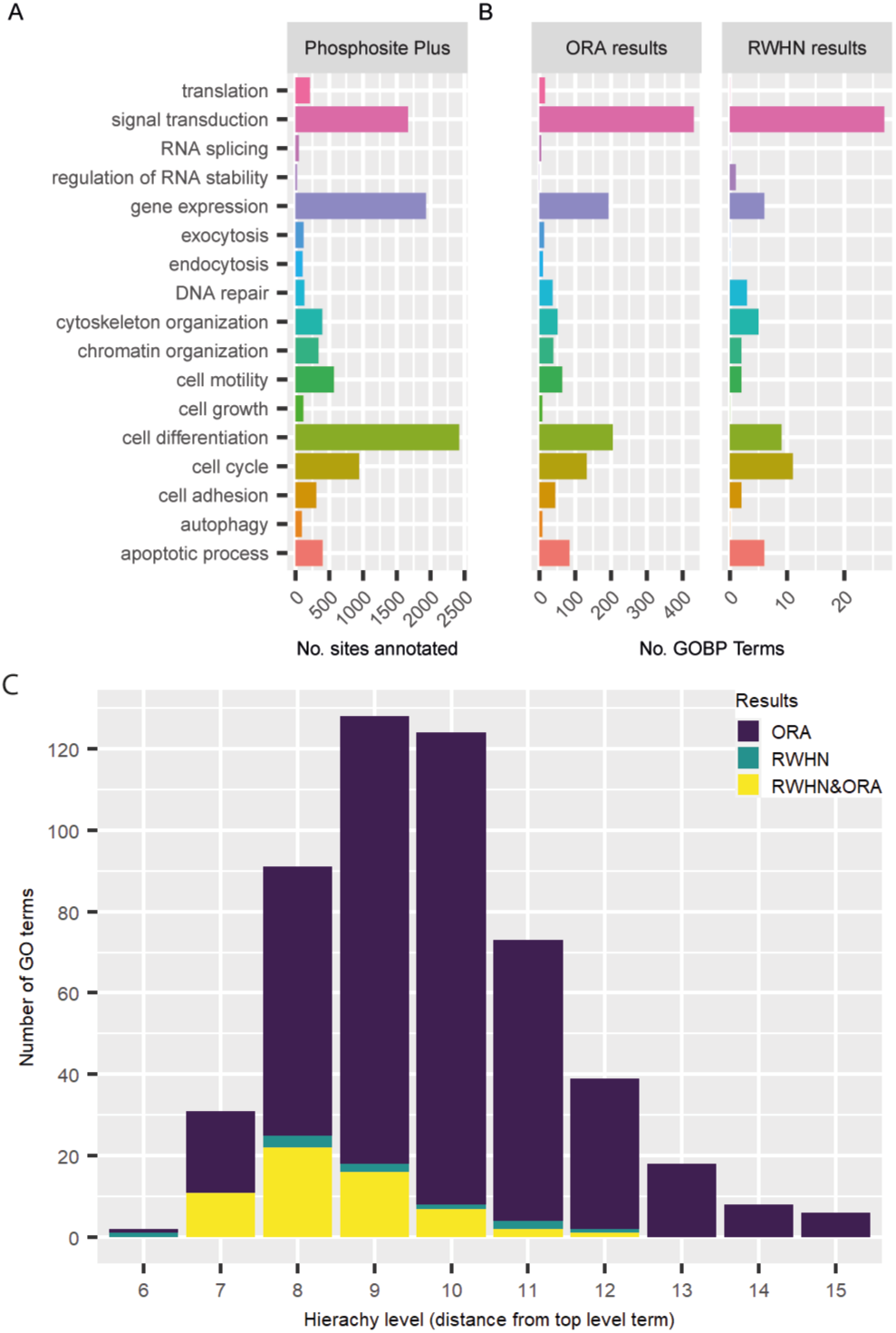
Distribution of annotations and classifications of sites extracted from PhosphoSitePlus. A) Distributions of functionally annotated sites (to mapped GO term). B) Distribution of terms in ORA results and RWHN results when analyzing functionally annotated sites from PSP. C) Number of terms from ORA or RWHN that are from different levels of the GO hierarchy

### Experimental Case Study 1: Dissecting EGF- and TGF-α-Induced Dynamic Phosphorylation

Given that PSP contains static annotations of sites from various sources, we wanted to test our algorithm on experimental datasets with controlled perturbations. We reasoned that this would allow us to assess whether our algorithm could differentially associate sites to their context-dependent function. The phosphoproteomics data retrieved from Francavilla *et al*. ^21^ describes the effect in HeLa cells of stimulation with EGF or TGF-α over a period of 90 minutes (with time points at 1, 8, 40 and 90 minutes). In this study, one of the findings was that EGF or TGF-α stimulation of EGFR induce receptor degradation or recycling, respectively. TGF-α was associated with a more potent mitogenic and migratory response than EGF over time ^21^. Further details on the experimental design and results from the publication can be found in Supplementary Table S2. Phosphorylated sites from Supplementary Table S1 of the original publication were filtered based on regulation by EGF or by TGF-α and divided based on which stimuli the regulation was dependent on. The separated data were clustered into 6 clusters using the fuzzy C-means method, as per the original publication, with each cluster representing a distinct dynamic profile of phosphorylation (Figure S3). A multilayer heterogeneous network was constructed for both sets of data, described in Table 2 and Table 3, respectively.

**Table 2.** Multilayer heterogeneous network, constructed from Francavilla *et al*. ^21^ EGF-regulated phosphorylated sites. GOBP terms were included as functional annotations.

| Subnetwork | Edges | Nodes |
| --- | --- | --- |
| Site-site | 4347 | 733 |
| Protein-protein | 16654 | 4072 |
| Function-function | 532 | 544 |
| Site-protein | 733 | 733 sites, 433 proteins |
| Protein-function | 1223 | 115 proteins, 532 functions |

**Table 3.** Multilayer heterogeneous network, constructed from Francavilla *et al*. ^21^ TGFα-regulated phosphorylated sites. GOBP terms were included as functional annotations.

| Subnetwork | Edges | Nodes |
| --- | --- | --- |
| Site-site | 3085 | 675 |
| Protein-protein | 16556 | 3960 |
| Function-function | 332 | 496 |
| Site-protein | 675 | 675 sites, 403 proteins |
| Protein-function | 1015 | 109 proteins, 486 functions |

Several biologically relevant terms were differentially ranked in the EGF and TGF-α networks (Figure 4A,C). Consistent with the original publication, the term “regulation of ERK1 and ERK2 cascade” was highly ranked using RWHN when seeds were set to sites in cluster 2 (representing EGF late responders) for the EGF network and in cluster 1 (representing TGF-α early responders) for the TGF-α network. Despite their central role to TGF-α signalling, when we performed standard ORA enrichment terms relating to MAPK/ERK were not found in any of the clusters (Figure 4D). However, both ORA and our method highlighted the role of the proteins belonging to TGF-α cluster 1 in regulating EGFR/ERBB signaling, with terms referring to regulation of these pathways ranked highly (Figure 4C,D). We used cluster 2 in the EGF condition and cluster 1 in the TGF-α condition to verify the robustness and accuracy of our approach. As the MAPK/ERK cascade is well studied in respect to EGF/TGF-α signaling, many of its associated sites have documented functions that we could use to benchmark our results. EGF cluster 2 contains sites of several known players of the MAPK/ERK cascades, including RSP6KA3_S375, SOS1 (T1119, S1137), RAF1 (S289, S96, S301), JUN_S243 and JUNB_259. All of these sites are predicted MAPK1/MAPK3 targets, with the exception of SOS1_T1119, which is a MAP2K1 target ^7^. As this cluster represents those sites phosphorylated late in the time course, we would expect ERK1/2 feedback regulation to emerge. Besides confirming the term “regulation of ERK1 and ERK2 cascade” in EGF cluster 2, results from the RHWN algorithm add detail to this picture, with “establishment of protein localization to organelle” ranked highly in this cluster (Figure 4A). This finding confirms the conclusion of Francavilla *et al*. of EGFR trafficking to the lysosome, and subsequent degradation, when stimulated by EGF ^21^. EGF cluster 2 is the only one which did not highly rank “regulation of clathrin-dependent endocytosis” compared to the 5 other clusters; as this cluster represents sites that are highly phosphorylated late in the time-course, this coincides with the understanding of receptor endocytosis as a process regulated in the early stage. This cluster is one of the five that highly ranks “regulation of cell migration” (Figure 4A); PSP also annotates several of the sites in this cluster to cell motility (PTPN12_S571, SLC9A1_S703, FLNA_S2152).

**Figure 4.**
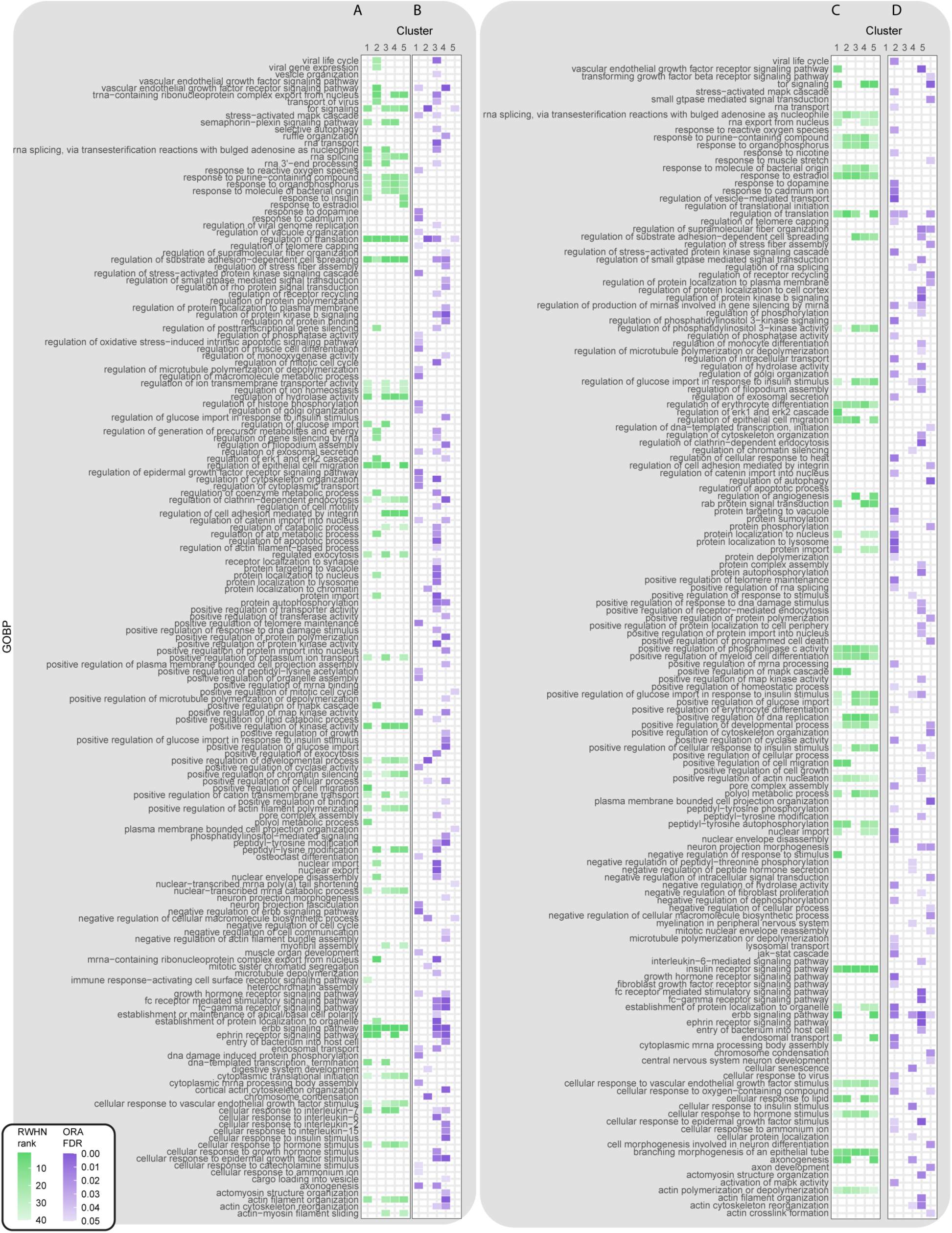
Site-centric and gene-centric analysis of Francavilla *et al*. ^21^ datasets. (A) RWHN results related to EGF regulated sites. (B) ORA results related to proteins modified upon EGF stimulation. (C) RWHN results related to TGF-α regulated sites. (D) ORA results related to proteins modified upon TGF-α stimulation.

TGF-α cluster 1, on the other hand, contains sites that are phosphorylated within the first eight minutes upon TGF-α stimulation. Accordingly, this cluster contains different members of the MAPK/ERK cascade, such as the canonical activating sites of MAPK1, MAPK3 and MAPK14 (also known as p38) and several sites on EGFR (EGFR_S991, EGFR_S995, EGFR_T693). EGFR_S991. A few of these sites, such as EGFR_T693, are annotated to “receptor internalization” in PSP. This cluster also contains several phosphorylated sites on known players in receptor trafficking, including RAB7A_Y183 and RAB11B_Y8 which are highlighted in the original publication ^21^, along with CBL_Y674 and SH3BP4_T118. Our algorithm ranked terms such as “endosomal transport,”, “establishment of protein localization to organelle” and “rab protein signal transduction” high when seed nodes were set to the sites found in this cluster (Figure 4C). This was only partially captured by the ORA (Figure 4D), which showed enrichment of the term “regulation of vesicle mediated transport” but also “protein localization to the lysosome”, the opposite of the experimentally derived conclusion of Francavilla *et al*.

EGF cluster 1 representing phosphorylated cycling sites ranks both “regulation of cell migration” and “positive regulation of cell migration” in the top 5% of terms but did not have a similar association with trafficking terms as TGF-α cluster 1 in the RWHN output. Consistent with Francavilla *et al*., we concluded that there was not an association between trafficking and regulation of migration upon EGF stimulation ^21^. Upon further investigation of this cluster, we found several sites that were shared with TGF-α cluster 1, such as the MAPK1/3/14 sites, EGFR_S991 and EGFR_T693. We also found the inhibitory RAF1 site S259 and SCRIB_1448. SCRIB has a putative role in cell migration ^45, 46^.

TGF-α clusters 4 and 5 also highly rank terms related to intracellular trafficking. In the ORA analysis (Figure 4D) cluster 4 was also enriched for “regulation of clathrin-dependent endocytosis”, corroborating this association. Of particular interest in cluster 4 are the EGFR (Y1110, Y1125 and Y1197), CBL (Y700) and CBLB (Y665. Y763. Y889) sites. None of the CBL/CBLB sites have roles described in PSP, but these proteins are known regulators of receptor in trafficking via clathrin ^47, 48^. Moreover, EGFR_Y1110 and EGFR_Y1197 are both annotated to receptor internalization in PSP, as well as inducing cell motility. This is in line with “regulation of epithelial cell migration” highly ranked in this cluster (Figure 4C). Taken together, this data indicates that TGF cluster 4 sites may have an initial role in regulating receptor internalization and migration, whilst the cluster 1 sites are regulating later parts of the process.

As a positive control for assignment of functions to clustered sites, sites annotated in PSP to the function “cell growth, inhibited” were added as a “spike-in” cluster to the EGF and TGF-α data, with sites already found in the data excluded from this new cluster. RWHN and ORA were then applied as previously. We found that both RWHN and ORA could assign these sites to the function “negative regulation of cell growth” (Figure 5A). However, ORA assigned 134 and 122 other terms to the sites added to the EGF and TGF data respectively, whilst RWHN assigned a refined list of 36 and 39 respectively (Figure 5B). This result suggests that RWHN assigns more specific functions to each phosphorylated site. Indeed, for each of the investigated clusters, RWHN was able to assign sites belonging to those clusters to functions that closely resemble those that have been experimentally proven by Francavilla *et al*. or described in PSP. The ORA results, whilst not inaccurate, assigned more generic biological functions and did not assign specific terms. These results suggest the RWHN algorithm can distinguish between sites of known function and can assign functional terms to differentially regulated phosphorylation sites that could be further investigated experimentally.

**Figure 5.**
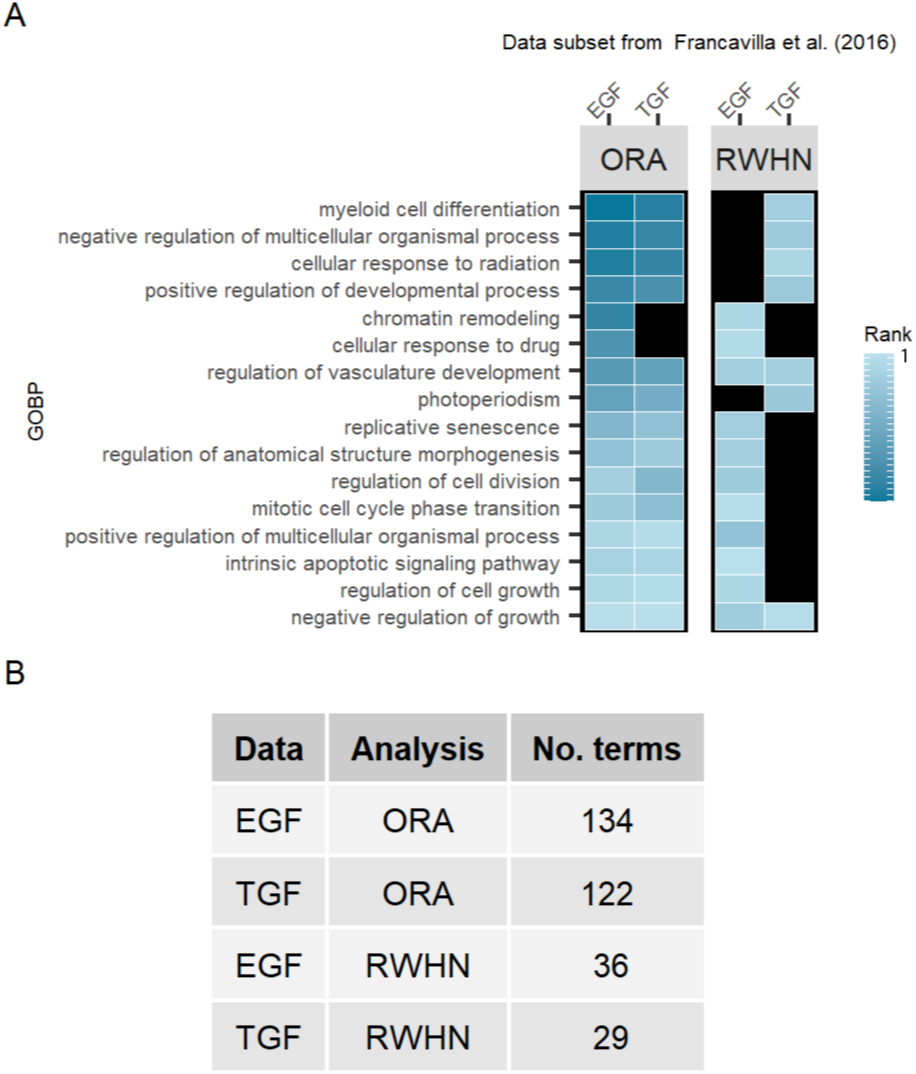
Cluster containing sites annotated to “Cell growth, inhibited” in Phosphosite Plus, spiked into Francavilla et al. networks. A) Terms in common between ORA and RWHN results with both the EGF and TGF-α networks. B) Total number of terms in the output of both types of analyses on both networks.

### Experimental Case study 2: Phosphorylation-dependent response to Lapatinib treatment and resistance in breast cancer

To verify the applicability of the algorithm with non-temporal shotgun phosphoproteomics data, we applied it to the experimental dataset from Ruprecht *et al.* ^22^ describing changes in the phosphoproteome upon treatment with the breast cancer drug lapatinib in sensitive (‘parental’ BT-474) or lapatinib-resistant (BT-474-J4) cells. The paper uncovered and experimentally validated the role of several metabolic enzymes and signaling pathways that were driving lapatinib resistance in breast cancer. In particular, proteins that form the spliceosome, those involved in glycolysis and glycogen catabolism and PI3K/AKT/mTOR pathway members were differentially phosphorylated in lapatinib-resistant cells compared to parental lapatinib-treated cells. Further details on the experimental design and results from the publication can be found in Supplementary Table S2.

We first constructed a multilayer heterogeneous network, as described in Experimental Procedures, which included 1603 phosphorylated sites that showed significant changes in response to lapatinib treatment in either the parental or resistant cell lines (Table 4). Data were grouped into 5 clusters (referred to as clusters 1 – 5) using the k-means method. The number of clusters to use was determined using the elbow plot method ^29^(Figure S4A), confirming that 5 clusters were sufficient to capture all features found in the data. Indeed, sites sharing similar regulation profiles between the resistant and parental cells were clustered together (Figure S4B). As the original publication used KEGG pathways rather than GO terms, we incorporated these in the functional-annotation layer of our network to investigate the flexibility of our approach.

**Table 4.** Multilayer heterogeneous network, constructed from Ruprecht *et al*. ^22^ lapatinib-regulated phosphorylated sites in parental and lapatinib-resistant cell lines. KEGG pathways were included as functional annotations.

| Subnetwork | Edges | Nodes |
| --- | --- | --- |
| Site-site | 75688 | 1599 |
| Protein-protein | 12752 | 3129 |
| Function-function | 15 | 152 |
| Site-protein | 1599 | 1599 sites, 930 proteins |
| Protein-function | 692 | 103 proteins, 152 pathways |

The results of RWHN applied to the multilayer heterogeneous network (Figure 6A) show that when the seed nodes are set to sites belonging to cluster 4, representing sites that are less phosphorylated in lapatinib-resistant cells, the spliceosome pathway was highly ranked. This cluster interestingly contained signalling sites such as ERBB2_Y1233, ERBB2_T1227, ERBB2_Y1233, ERBB3_S627 and MAPK1_Y187, alongside sites on known spliceosome factors such as SRRM2 (S454, S2449), SRSF6 (S301) CWC25 (S170). Figure 6B shows that there was no enrichment for any pathways in this cluster using standard ORA. However, ORA did show enrichment for metabolic terms in several clusters, unlike RWHN, despite this being one of the clear conclusions of the original paper. Metabolic pathways were enriched in clusters 3 (“glycolysis / gluconeogenesis”) and 5 (“central carbon metabolism in cancer”) using ORA. Since much of the investigation and experimental validation by Ruprecht *et al*. ^22^ was done on the sites found to be regulated only in lapatinib-treated resistant cells, we investigated whether our approach could uncover more nuanced roles of sites within these datasets. We constructed a second network using only those sites that were regulated in lapatinib-treated resistant cells (Table 5). These were grouped into four clusters (referred to as clusters 1 – 4) by K-means clustering as before (Figure S4A,C) and KEGG pathways were used again in the functional annotation layer.

**Figure 6.**
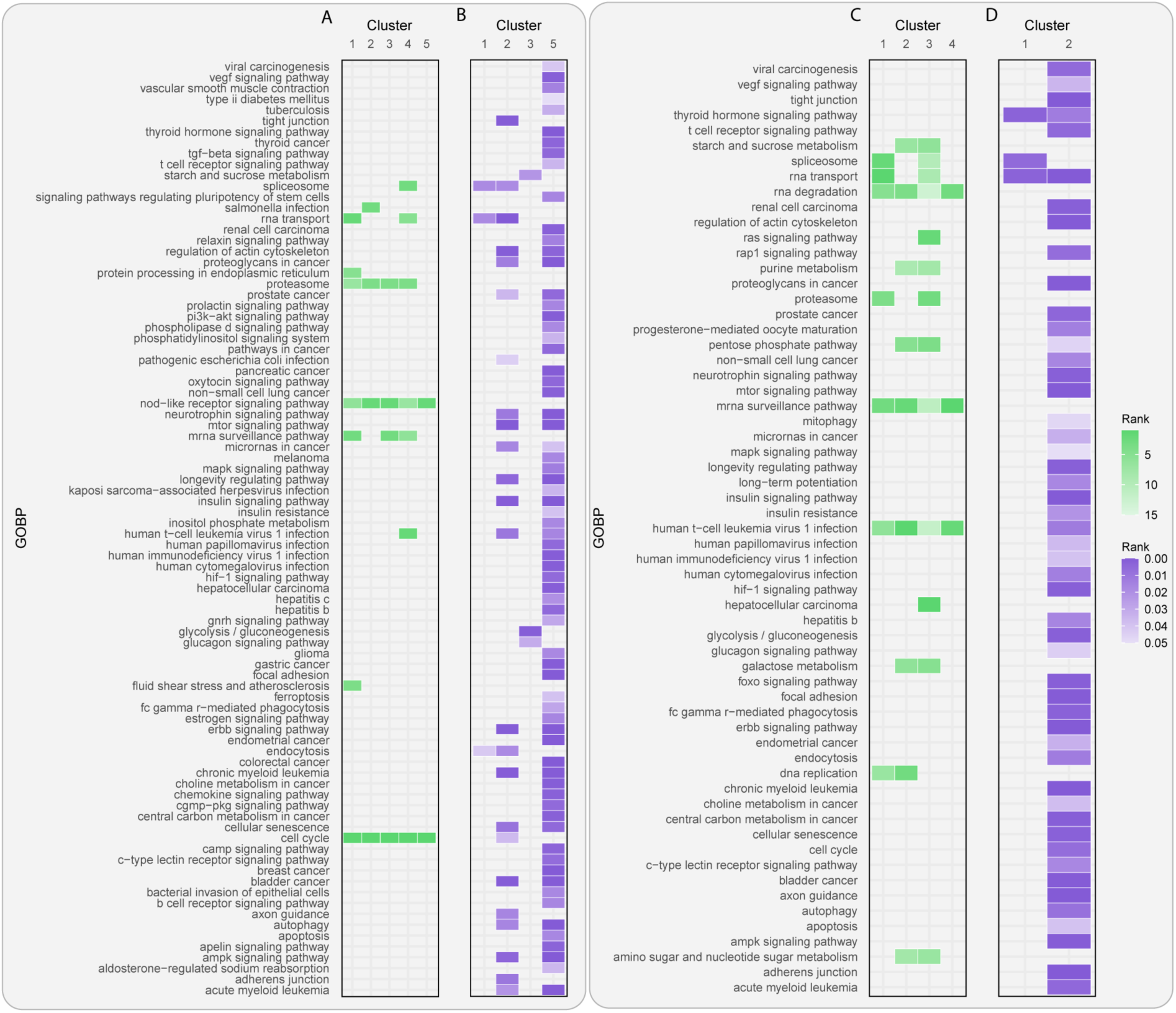
Site-centric and gene-centric analysis of Ruprecht *et al*. ^22^ datasets. (A) RWHN results related to sites regulated by lapatinib in lapatinib-resistant and parental cell lines. (B) ORA results related to proteins modified by lapatinib in lapatinib-resistant and parental cell lines. (C) RWHN results related to sites regulated by lapatinib in lapatinib-resistant cell lines only. (D) ORA results related to proteins modified by lapatinib in lapatinib-resistant cell lines only.

**Table 5.**
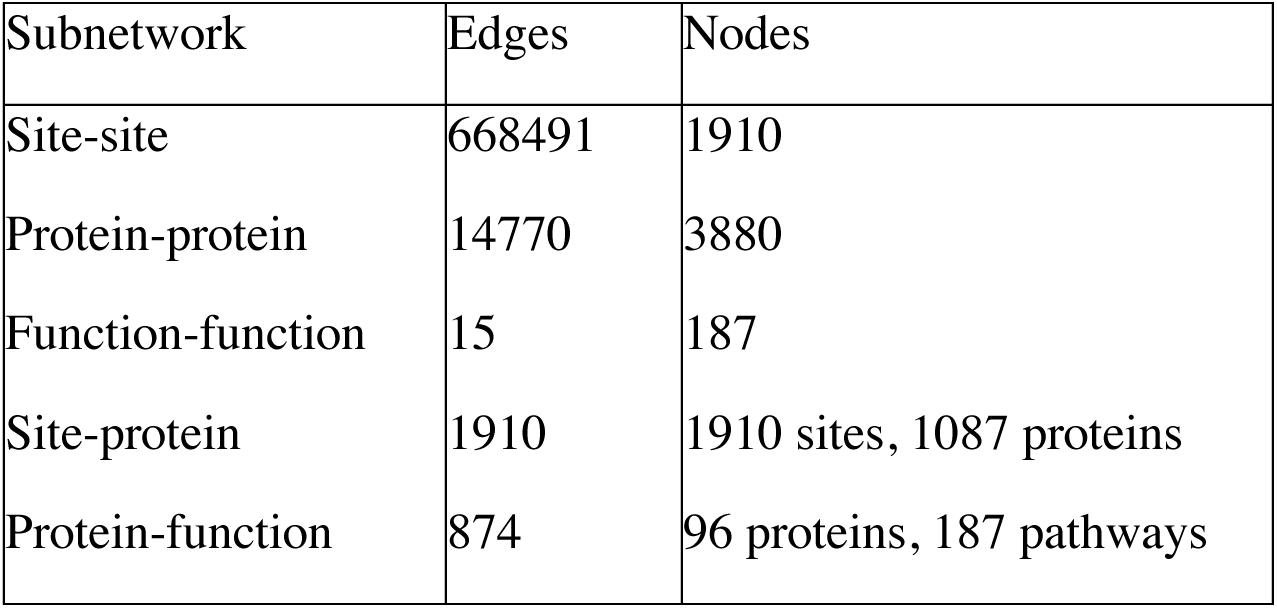
Multilayer heterogeneous network, constructed from Ruprecht *et al*. ^22^ lapatinib-regulated phosphorylated sites in lapatinib-resistant cell line. KEGG pathways were included as functional annotations.

As described in Figure 6C, two sets of seed nodes (clusters 2 and 3, representing sites substantially upregulated and downregulated, Figure S4C) tended to rank metabolic pathways more highly (“starch and sucrose metabolism”, “galactose metabolism”, “pentose phosphate pathway”, “purine metabolism”, “amino sugar and nucleotide sugar metabolism”). Cluster 3, along with cluster 1 (representing moderately downregulated sites), ranked “spliceosome” highly. These initial interpretations indicate that using a specific subset of the regulated sites in the data yielded more specific and functionally relevant terms being ranked highly by the RWHN algorithm. Moreover, only cluster 2 showed enrichment for any metabolic pathways using ORA (Figure 6D). As this is a large cluster containing all the sites shown to be significantly upregulated in the resistant cell line, there is a considerable amount of ‘noise’ in the ORA results. The enrichment of two metabolic pathways (“glycolysis / gluconeogenesis” and “central carbon metabolism in cancer”) potentially corroborate the role of these sites as metabolism regulators, as indicated by RWHN. This cluster contained several of the sites classified as metabolic drivers of the resistance phenotype, such as ENO1_Y44, PKM_Y175, PDHA_S293/S300. Intriguingly, S17 on the diverse signalling regulator SRC is also found in this cluster; this site is annotated to induction of enzymatic activity in PSP.

A closer look at cluster 3 uncovered sites on spliceosome factors (RBMX_S58, CDK13_S525, CWC25_S17) and known signaling proteins (ERBB2_Y1233, FGFR4_S505). A site on the adaptor protein IRS2 (S1176) was also found in this cluster; as a known player in glucose and lipid metabolism ^49^. This points towards the link between rewired signalling and metabolism investigated by Ruprecht *et al*. ^22^. It is also annotated to “metastatic potential” in PSP, indicating its therapeutic potential. ORA also showed enrichment for “spliceosome” in cluster 1. Indeed, this cluster contained several of the sites highlighted in the paper. For example, the known lapatinib responsive sites SF3B2_S309 and SF3B1_T261 could be found in this cluster, alongside 17 of the 30 SRRM2 sites that were significantly regulated by lapatinib.

From the comparison of ORA and the RWHN method we have extracted meaningful associations between specific phosphorylated sites and their functions in driving lapatinib resistance. Our algorithm also highlighted potential further sites, such as IRS2_S1176, compared to the ORA method, that may serve as hubs or intersections of crosstalk between key signalling pathways and the rewired metabolic processes. It can be noted that the RWHN output was substantially more refined than the ORA output, with the conclusions of the original paper being captured in 14 terms in our analysis of the sites regulated in resistance, as opposed to 51 using ORA.

### Experimental Case Study 3: Spatially resolved activity of HRAS

The final dataset we analyzed described immunoprecipitation (IP) samples, in order for us to see if our algorithm would be useful for analyzing local as well as global phosphoproteomes. The subcellular localization of signaling proteins has been established as crucial in regulating cascades and cellular processes. There are now numerous methods that allow experimental biologists to detect spatial effects of proteins and phosphorylation, for example through proximity labeling and targeted proteomics^50, 51^. Recent work by Santra *et al*. ^23^ took a similar approach to investigate the differential roles of a mutant, constitutively active form of HRAS, HRASV12, at different subcellular locations. HRASV12 constructs tagged with different signal peptides (targeting them to different subcellular locations) were stably transfected into HeLa cells; this allowed the authors to collect multi-omic samples with endogenous HRAS, unlocalized HRASV12 and HRASV12 localized to either the plasma membrane in disordered membrane regions (DM) or lipid rafts (LR), Golgi apparatus (GA) and endoplasmic reticulum (ER). Using this approach, it was confirmed that the majority of HRASV12 activity is mediated from the plasma membrane, with ER- and GA-localized HRASV12 involved in regulating some of HRASV12’s role in cell survival. Further details on the experimental design and results from the publication can be found in Supplementary Table S2.

The phosphoproteomics data from this study were clustered using k-means clustering, with an optimal cluster number of 6 determined using an elbow plot (Figure S5A,B). Table 6 describes the properties of the multilayer network constructed with these clusters. The first three clusters represented dynamics discussed in the original paper: cluster 1 represented phosphorylated events of mutant HRAS regardless of localization (location-independent); cluster 4 represented HRAS in the secretory pathway, at the ER and GA (ER/GA); cluster 6 represented HRAS at the plasma membrane, in the DM or LR (DM/LR). As these experimental conditions were best characterized in the original paper, downstream analysis focused on these three clusters.

**Table 6.** Multilayer heterogeneous network, constructed using regulated sites from the Santra *et al.* ^23^ dataset. GOBP terms were included as functional annotations.

| Subnetwork | Edges | Nodes |
| --- | --- | --- |
| Site-site | 766 | 621 |
| Protein-protein | 7262 | 2163 |
| Function-function | 370 | 452 |
| Site-protein | 621 | 621 sites, 474 proteins |
| Protein-function | 900 | 122 proteins, 442 functions |

Using RWHN (Figure 7, in purple), we found that the ER/GA and DM/LR clusters both highly ranked key signaling pathways such as “ERK1 and ERK2 cascade” and “TOR signaling”, alongside signaling regulation such as “Receptor mediated endocytosis”. This is consistent with the finding from the original paper, that the majority of HRAS mediated effects derive from the localized population. Specifically, the DM/LR cluster ranks mitogenic processes highly, such as “regulation of epithelial cell proliferation” and “positive regulation of metaphase/anaphase transition of cell cycle”. The original paper also concluded that the majority of HRAS’ mitogenic effects were mediated from the plasma membrane. The ER/GA cluster independently ranks “positive regulation of growth” and “Notch signaling pathway”. This potentially implicates ER/GA localized HRAS, in the known developmental roles of HRAS in development; this was touched upon but not investigated in the original paper.

**Figure 7.**
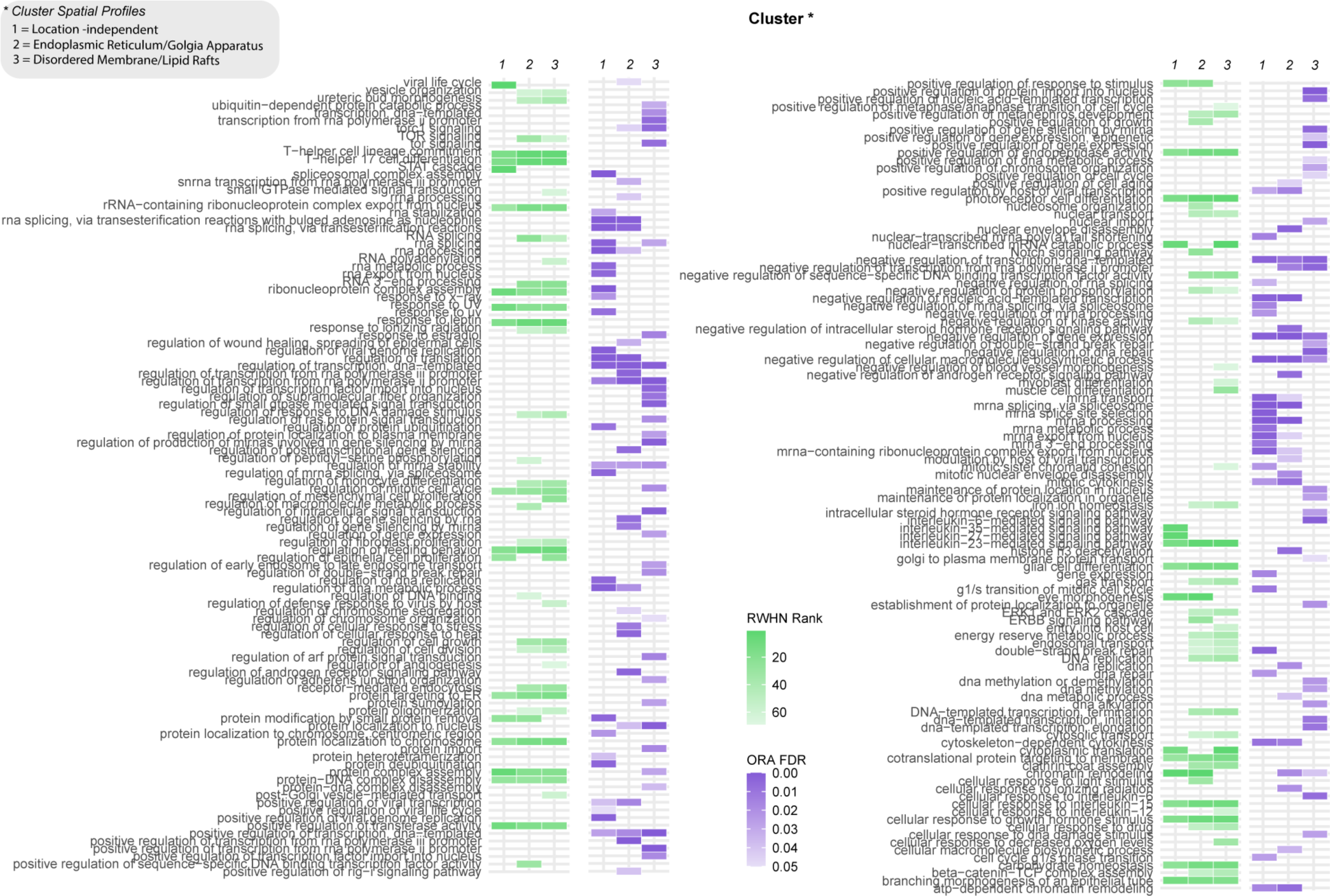
Site-centric and gene-centric analysis of Santra *et al.* ^23^ datasets; results from RWHN and ORA are in green and purple, respectively. Clusters shown in the figure refer to phosphorylated sites that are location-independent or found at the endoplasmic reticulum (ER)/Golgi apparatus (GA) or disordered membrane (DM)/ lipid rafts (LR) (marked in the figure as clusters 1, 2, and 3, respectively).

Conversely, where a term is ranked highly in all three clusters, this could indicate that this is part of the localization-independent regulation of HRAS. Twenty of the highly regulated terms fit this definition; of these, several are related to immunological processes, such as T-help cell lineage commitment” and “cellular response to interleukin-25”. The location-independent cluster also independently ranks “interleukin-27-mediated signaling pathway” and “STAT cascade”. This suggests that HRAS’ role in regulating cytokine and immune response is not location dependent.

An obvious outcome of the results using RWHN and ORA on this data is the difference in the number terms associated to each cluster. There were 132 terms describing these clusters with ORA (Figure 7, in green), compared to 86 with RWHN, with many of the enriched terms related to the overrepresented categories depicted in Figure 3, such as “gene expression”. Several comparisons can be drawn between the results of the two analyses to add credence to the accuracy of results from RWHN. For example, cluster 1 shows enrichment for the term “Interleukin-6-mediated signaling pathway”, and cluster 3 is enriched for the term “regulation of mitotic cell cycle”. These results from ORA are consistent with the results from RWHN and the original paper, but with a significant amount of noise and redundancy in the ORA output relative to RWHN.

## 5. Discussion

Phosphorylation has an important impact on protein function and thus cellular behavior. Within a network of kinase-substrate interactions, phosphorylation modulates the flow of information and regulation of disparate processes throughout the cell. Understanding the impact of phosphorylation on protein function traditionally required the experimental investigation and manipulation of individual sites, however with the advent of high throughput phosphoproteomics thousands of novel phosphorylation sites of unknown function have been discovered. As there is limited experimentally validated information available on how regulation at individual phosphorylated sites impacts the downstream cellular output, current methods for analyzing high throughput phospho-proteomics data rely on general descriptors of phosphorylated protein function such as GO terms or involvement in known pathways. Their use in proteomics and transcriptomics analysis, though widespread, has come under recent criticism. Maertens *et al* ^52^ demonstrated that annotations are lacking for a substantial number of genes known to be important in cancer, whilst Haynes *et al*. ^53^ demonstrate a “rich-getting-richer” effect, with well-studied genes having higher annotation rates despite having less-omic or molecular evidence for the association with processes or diseases. These problems are exacerbated when used in analyzing phosphoproteomics data, as enrichment analyses also disregard site-specific or multiple-site regulation, masking the roles of proteins in a particular modification state ^10, 12^.

Here, we developed and tested a method that associates phosphorylated sites to potential function using RWHN. By incorporating the pattern of phosphorylation upon perturbation, we consider more of the information available in phosphoproteomics datasets and establish the phosphorylated sites as the key drivers in the functional analysis. The algorithm is capable of recapturing experimentally proven functions of phosphorylated sites in a non-gene centric manner, refining analysis and extracting more information from phosphoproteomics data.

To prove the utility of our algorithm we applied it to a validation network, which represented a simple model of phosphorylation dynamics in the MAPK/ERK pathway, and to two previously published phosphoproteomics dataset describing temporal cellular signalling events and the impact of resistance to drug treatment in breast cancer, respectively. Our algorithm successfully distinguished between differentially regulated phosphorylated sites from the same protein and associated them to both known and previously uninvestigated functions. For example, when applying this approach to the data from Francavilla *et al.* ^21^, we can recapture the roles of EGFR sites with functions described in PSP. Both EGFR_Y1110 and EGFR_Y1197 are annotated to receptor internalization and cell motility in PSP; in our analysis, we found that these sites are both found in clusters that associate with “regulation of clathrin dependent endocytosis”, however only the TGF-α cluster also ranked “regulation of cell migration” highly, reproducing the experimental evidence from Francavilla *et al*. Whilst the authors of the original manuscript associated these sites with their functions using a combination of gene-centric approaches and experimental validation, here we use a single site-centric algorithm to extract these associations directly from the phosphoproteomics dataset, demonstrating the power of our approach in quickly and more specifically narrowing down candidates for further functional studies. By associating ligand-dependent regulation of phosphorylated sites, it is possible to disentangle their multifaceted role in regulating cellular signalling networks and downstream cellular behaviors.

From the Ruprecht *et al*. data, we can use the example of SRRM2 to illustrate the validity of our approach. This component of the spliceosome is known to be highly phosphorylated, with 675 phosphorylation sites recorded in PSP. At the time of writing none of these sites was ascribed to a function in the database, despite the importance of phosphorylation and dephosphorylation events in orchestrating splicing events ^54^. It is also regularly mutated (>5% of cases in TGCA) in lung, stomach, bladder, endometrial and colorectal cancers ^6^. Ruprecht *et al*. uncovered a new role for the spliceosome in modulating lapatinib-resistance, highlighting several sites on spliceosome proteins (including SRRM2_S1132, S1987 and S970) that were differentially modified in resistant and parental cell populations. Previous work has described how SRRM2-depletion in HER2-positive breast and ovarian cancer cells reduced the rate of migration ^55^; the spliceosome plays myriad roles in the breast cancer environment, but site-specific analysis is lacking. Using our algorithm, we can begin to uncover the nuanced modification-specific roles of proteins such as SRRM2 in metabolic rewiring. We suggest that eleven of the SRRM2 sites found in cluster 2 of the lapatinib-resistant sites (Figure 6C) could be at the intersection between the spliceosome and metabolic processes; the conjunction between these processes was central to the work of Ruprecht *et al*. The sites in this cluster could be investigated as points of crosstalk between these different cellular processes. This example highlights the value of our approach in generating hypotheses of modified protein function and signaling pathway crosstalk.

Although the associations predicted here between sites and functions cannot be interpreted as causative, phosphorylation sites can act as indirect, and context-dependent, regulators in cellular processes. This is demonstrated by the clustering of SRC_S17, a protein that is typically associated with signalling downstream of cell-surface receptors, with a cluster that ranked metabolic pathways higher than cellular processing or signalling pathways. Critical to this is a robust method of defining clusters, to capture meaningful regulatory patterns at different phosphorylated sites. Here we define clusters using established techniques (e.g., elbow method for K-means) or visually identifying those with differing time-resolved behaviors. By using clusters to define edges between phosphorylated sites in the multilayer heterogeneous network, the pattern of experimentally perturbed modules in the cellular system can be investigated. Although having fixed edges between sites, as with the PPI layer of the multilayer network, would likely allow for a closer representation of the whole cellular system, this would not be computationally viable nor in the interest of experimental biologists. This algorithm is targeted towards datasets that describe the effect of specific treatments or perturbations, allowing for a focused view on the impacted processes to identify candidates to investigate further. Quantitatively defining the edges between phosphorylated sites based on the dataset collected also prevents the issues plaguing standard enrichments analyses as discussed before; fixed edges, defined on information in databases or previously published data, would almost certainly skew the analysis towards well studied phosphorylated sites and biological processes.

Multilayer heterogeneous networks are increasingly being used to integrate ‘omic data; here, their use allows phosphoproteomics data to be incorporated with PPI networks and functional annotations, overcoming the issue of considering phosphoproteomics data primarily on the protein level. A potential drawback of our approach is the reliance on large semi-curated resources such GO or STRING. For instance, different clusters may have many terms or pathways in common given the involvement of many proteins in the same biological functions and the high proportion of frequently used GO terms ^56^. We theorize that this may be rectified if the data were clustered into more groups, in order to capture more nuanced phosphorylation patterns. However, enrichment of non-specific terms or false positives remains an issue when analyzing high-throughput ‘omics data by other commonly used methods too. By only including edges in the protein layer of the network with a STRING experimental confidence score of greater than 0.4 and only including non-redundant terms in the function layer we have reduced the noise in the resulting list of associated terms. This could be taken further by filtering for only high confidence functional annotations or incorporating annotations from multiple sources into the network. Moreover, RWHN does not result in a different set of terms in the output compared to the accepted standard method ORA, but rather provides a refinement that will allow for biological insights to be reached more clearly.

Our method is flexible enough to be used with any discovery phosphoproteomics data that describes a change between conditions. This is an improvement on the previously published use of RWHN using multiple sources of phosphoproteomics data to uncover disease-dependent regulation ^19^. Moreover, it could easily be generalized to any post-translational modification proteomics dataset, as it incorporates readily available PPI and functional annotation data, as demonstrated here. The fundamental aspect would be maintained with any of these expansions: specific patterns of regulation at modified sites dictate movement through the multilayer heterogeneous network.

## 6. Conclusion

We have proposed a site-centric approach to analyze phosphoproteomics data, that provides a robust alternative to gene-centric methods of analysis. We integrated clustered quantitative phosphoproteomics data, a context-specific PPI network and functional annotations into a multilayer, heterogeneous network and used the RWHN method to predict the functions of phosphorylation sites with similar regulatory patterns. Using our algorithm, we extracted experimentally validated associations between phosphorylated sites and their role in cellular processes, which could not be captured using the typical gene-centric methods. Moreover, our algorithm has the potential to be used by researchers in predicting novel site-function associations and generating hypotheses to be experimentally validated.

## 7. Supporting Information

Figure S1. Optimization of RWHN parameters using the validation dataset.

Figure S2. Validation using randomly permuted networks.

Figure S3. Clustering of Francavilla et al. data.

Figure S4. Clustering of Ruprecht et al. data.

Figure S5. Clustering of Santra et al. data.

Table S1. PSP ON_PROCESS annotations, that describe biological function of sites, mapped to most closely related GOBP term.

Table S2. Summary of the experimental methods and results from each of the Case Study datasets.

Data and source code for the analysis described in the manuscript can be found at github.com/JoWatson2011/RWHN_Phosphoproteomics

An R package for use of the tool can be found at github.com/JoWatson2011/phosphoRWHN

## 8. Author Contributions

The manuscript was written through contributions of all authors. All authors have given approval to the final version of the manuscript.

## 10. Acknowledgements

We would like to acknowledge the assistance given by Research IT and the use of the Computational Shared Facility at The University of Manchester. We would like to thank current and former members of the Schwartz and Francavilla labs for their thoughtful and insightful discussion on this work and Dr Joe Swift (University of Manchester) for useful comments on our manuscript.

## 11. Abbreviations

RWHN: Random walk on heterogeneous network
PTM: Post-translational modification
PPI: Protein-protein interaction
GO: Gene Ontology
PSP: Phosphosite Plus
EGF: Epidermal Growth Factor
TGF-α: Tumor growth factor α
RWR: Random walk with restart
GOBP: Gene Ontology Biological Processes
FDR: False discovery rate
SILAC: Stable Isotope Labelling of Amino Acids in Cell culture
KDE: Kernel Density Estimation

## References

(1) Sugiyama, N.; Imamura, H.; Ishihama, Y. Large-Scale Discovery of Substrates of the Human Kinome. Scientific Reports 2019, 9 (1), 10503. https://doi.org/10.1038/s41598-019-46385-4.

(2) Day, E. K.; Sosale, N. G.; Lazzara, M. J. Cell Signaling Regulation by Protein Phosphorylation: A Multivariate, Heterogeneous, and Context-Dependent Process. Current Opinion in Biotechnology 2016, 40, 185–192. https://doi.org/10.1016/j.copbio.2016.06.005.

(3) Vyse, S.; Desmond, H.; Huang, P. H. Advances in Mass Spectrometry Based Strategies to Study Receptor Tyrosine Kinases. IUCrJ 2017, 4 (Pt 2), 119–130. https://doi.org/10.1107/S2052252516020546.

(4) Munk, S.; Refsgaard, J. C.; Olsen, J. V. Systems Analysis for Interpretation of Phosphoproteomics Data. In Phospho-Proteomics: Methods and Protocols; von Stechow, L., Ed.; Springer New York: New York, NY, 2016; pp 341–360. https://doi.org/10.1007/978-1-4939-3049-4_23.

(5) Sacco, F.; Perfetto, L.; Cesareni, G. Combining Phosphoproteomics Datasets and Literature Information to Reveal the Functional Connections in a Cell Phosphorylation Network. PROTEOMICS 2018, 18 (5–6), 1700311. https://doi.org/10.1002/pmic.201700311.

(6) Hornbeck, P. V.; Zhang, B.; Murray, B.; Kornhauser, J. M.; Latham, V.; Skrzypek, E. PhosphoSitePlus, 2014: Mutations, PTMs and Recalibrations. Nucleic Acids Research 2015, 43 (D1), D512–D520. https://doi.org/10.1093/nar/gku1267.

(7) Linding, R.; Jensen, L. J.; Pasculescu, A.; Olhovsky, M.; Colwill, K.; Bork, P.; Yaffe, M. B.; Pawson, T. NetworKIN: A Resource for Exploring Cellular Phosphorylation Networks. Nucleic Acids Research 2008. https://doi.org/10.1093/nar/gkm902.

(8) Casado, P.; Rodriguez-Prados, J.-C.; Cosulich, S. C.; Guichard, S.; Vanhaesebroeck, B.; Joel, S.; Cutillas, P. R. Kinase-Substrate Enrichment Analysis Provides Insights into the Heterogeneity of Signaling Pathway Activation in Leukemia Cells. Science signaling 2013, 6 (268), rs6. https://doi.org/10.1126/scisignal.2003573.

(9) Mischnik, M.; Sacco, F.; Cox, J.; Schneider, H.-C.; Schäfer, M.; Hendlich, M.; Crowther, D.; Mann, M.; Klabunde, T. IKAP: A Heuristic Framework for Inference of Kinase Activities from Phosphoproteomics Data. Bioinformatics 2016, 32 (3), 424–431. https://doi.org/10.1093/bioinformatics/btv699.

(10) Needham, E. J.; Parker, B. L.; Burykin, T.; James, D. E.; Humphrey, S. J. Illuminating the Dark Phosphoproteome. Science Signaling 2019, 12 (565), eaau8645. https://doi.org/10.1126/scisignal.aau8645.

(11) Rudolph, J. D.; de Graauw, M.; van de Water, B.; Geiger, T.; Sharan, R. Elucidation of Signaling Pathways from Large-Scale Phosphoproteomic Data Using Protein Interaction Networks. Cell systems 2016, 3 (6), 585–593.e3. https://doi.org/10.1016/j.cels.2016.11.005.

(12) Krug, K.; Mertins, P.; Zhang, B.; Hornbeck, P.; Raju, R.; Ahmad, R.; Szucs, M.; Mundt, F.; Forestier, D.; Jane-Valbuena, J.; Keshishian, H.; Gillette, M. A.; Tamayo, P.; Mesirov, P.; Jaffe, J. D.; Carr, S. A.; Mani, D. R. A Curated Resource for Phosphosite-Specific Signature Analysis. Molecular & cellular proteomics : MCP 2019, 18 (3), 576–593. https://doi.org/10.1074/mcp.TIR118.000943.

(13) Boyle, E. I.; Weng, S.; Gollub, J.; Jin, H.; Botstein, D.; Cherry, J. M.; Sherlock, G. GO::TermFinder—Open Source Software for Accessing Gene Ontology Information and Finding Significantly Enriched Gene Ontology Terms Associated with a List of Genes. Bioinformatics 2004, 20 (18), 3710–3715. https://doi.org/10.1093/bioinformatics/bth456.

(14) Battiston, F.; Nicosia, V.; Chavez, M.; Latora, V. Multilayer Motif Analysis of Brain Networks. Chaos: An Interdisciplinary Journal of Nonlinear Science 2017, 27 (4), 047404. https://doi.org/10.1063/1.4979282.

(15) Li, Y.; Patra, J. C. Genome-Wide Inferring Gene–Phenotype Relationship by Walking on the Heterogeneous Network. Bioinformatics 2010, 26 (9), 1219–1224. https://doi.org/10.1093/bioinformatics/btq108.

(16) Jiang, R. Walking on Multiple Disease-Gene Networks to Prioritize Candidate Genes. JOURNAL OF MOLECULAR CELL BIOLOGY 2015, 7 (3, SI), 214–230. https://doi.org/10.1093/jmcb/mjv008.

(17) Soul, J.; Hardingham, T. E.; Boot-Handford, R. P.; Schwartz, J.-M. PhenomeExpress: A Refined Network Analysis of Expression Datasets by Inclusion of Known Disease Phenotypes. Scientific Reports 2015, 5 (1), 8117. https://doi.org/10.1038/srep08117.

(18) Tang, Y.; Chen, K.; Wu, X.; Wei, Z.; Zhang, S.-Y.; Song, B.; Zhang, S.-W.; Huang, Y.; Meng, J. DRUM: Inference of Disease-Associated M6A RNA Methylation Sites From a Multi-Layer Heterogeneous Network. Frontiers in Genetics 2019, 10, 266. https://doi.org/10.3389/fgene.2019.00266.

(19) Xu, X.; Wang, M. Inferring Disease Associated Phosphorylation Sites via Random Walk on Multi-Layer Heterogeneous Network. IEEE/ACM Transactions on Computational Biology and Bioinformatics 2016, 13 (5), 836–844. https://doi.org/10.1109/TCBB.2015.2498548.

(20) Behar, M.; Hoffmann, A. Understanding the Temporal Codes of Intra-Cellular Signals. Current Opinion in Genetics & Development 2010, 20 (6), 684–693. https://doi.org/10.1016/J.GDE.2010.09.007.

(21) Francavilla, C.; Papetti, M.; Rigbolt, K. T. G.; Pedersen, A.-K.; Sigurdsson, J. O.; Cazzamali, G.; Karemore, G.; Blagoev, B.; Olsen, J. V. Multilayered Proteomics Reveals Molecular Switches Dictating Ligand-Dependent EGFR Trafficking. Nature Structural & Molecular Biology 2016, 23 (6), 608–618. https://doi.org/10.1038/nsmb.3218.

(22) Ruprecht, B.; Zaal, E. A.; Zecha, J.; Wu, W.; Berkers, C. R.; Kuster, B.; Lemeer, S. Lapatinib Resistance in Breast Cancer Cells Is Accompanied by Phosphorylation-Mediated Reprogramming of Glycolysis. Cancer Research 2017, 77 (8), 1842–1853. https://doi.org/10.1158/0008-5472.CAN-16-2976.

(23) Santra, T.; Herrero, A.; Rodriguez, J.; von Kriegsheim, A.; Iglesias-Martinez, L. F.; Schwarzl, T.; Higgins, D.; Aye, T.-T.; Heck, A. J. R.; Calvo, F.; Agudo-Ibáñez, L.; Crespo, P.; Matallanas, D.; Kolch, W. An Integrated Global Analysis of Compartmentalized HRAS Signaling. Cell reports 2019, 26 (11), 3100–3115.e7. https://doi.org/10.1016/j.celrep.2019.02.038.

(24) Cox, J.; Mann, M. MaxQuant Enables High Peptide Identification Rates, Individualized p.p.b.-Range Mass Accuracies and Proteome-Wide Protein Quantification. Nature Biotechnology 2008, 26 (12), 1367–1372. https://doi.org/10.1038/nbt.1511.

(25) J. Tyanova, S.; Temu, T.; Sinitcyn, P.; Carlson, A.; Hein, M. Y.; Geiger, T.; Mann, M.; Cox, The Perseus Computational Platform for Comprehensive Analysis of (Prote)Omics Data. Nature Methods 2016, 13 (9), 731–740. https://doi.org/10.1038/nmeth.3901.

(26) Ritchie, M. E.; Phipson, B.; Wu, D.; Hu, Y.; Law, C. W.; Shi, W.; Smyth, G. K. Limma Powers Differential Expression Analyses for RNA-Sequencing and Microarray Studies. Nucleic Acids Research 2015, 43 (7), e47–e47. https://doi.org/10.1093/nar/gkv007.

(27) Schwämmle, V.; Jensen, O. N. A Simple and Fast Method to Determine the Parameters for Fuzzy c–Means Cluster Analysis. Bioinformatics 2010, 26 (22), 2841–2848. https://doi.org/10.1093/bioinformatics/btq534.

(28) Campello, R. J. G. B.; Hruschka, E. R. A Fuzzy Extension of the Silhouette Width Criterion for Cluster Analysis. Fuzzy Sets and Systems 2006, 157 (21), 2858–2875. https://doi.org/10.1016/J.FSS.2006.07.006.

(29) Zambelli, A. E. A Data-Driven Approach to Estimating the Number of Clusters in Hierarchical Clustering. F1000Research 2016, 5, 2809. https://doi.org/10.12688/f1000re-search.10103.1.

(30) Szklarczyk, D.; Gable, A. L.; Lyon, D.; Junge, A.; Wyder, S.; Huerta-Cepas, J.; Simonovic, M.; Doncheva, N. T.; Morris, J. H.; Bork, P.; Jensen, L. J.; Mering, C. von. STRING V11: Protein–Protein Association Networks with Increased Coverage, Supporting Functional Discovery in Genome-Wide Experimental Datasets. Nucleic Acids Research 2018, 47 (D1), D607–D613. https://doi.org/10.1093/nar/gky1131.

(31) Yu, G.; Li, F.; Qin, Y.; Bo, X.; Wu, Y.; Wang, S. GOSemSim: An R Package for Measuring Semantic Similarity among GO Terms and Gene Products. Bioinformatics 2010, 26 (7), 976–978. https://doi.org/10.1093/bioinformatics/btq064.

(32) Wang, J. Z.; Du, Z.; Payattakool, R.; Yu, P. S.; Chen, C.-F. A New Method to Measure the Semantic Similarity of GO Terms. Bioinformatics 2007, 23 (10), 1274–1281. https://doi.org/10.1093/bioinformatics/btm087.

(33) Stoney, R. A.; Ames, R. M.; Nenadic, G.; Robertson, D. L.; Schwartz, J.-M. Disentangling the Multigenic and Pleiotropic Nature of Molecular Function. BMC systems biology 2015, 9 Suppl 6 (Suppl 6), S3. https://doi.org/10.1186/1752-0509-9-S6-S3.

(34) Blondel, V. D.; Guillaume, J.-L.; Lambiotte, R.; Lefebvre, E. Fast Unfolding of Communities in Large Networks. J. Stat. Mech. 2008, 2008 (10), P10008. https://doi.org/10.1088/1742-5468/2008/10/P10008.

(35) Kuleshov, M. V; Jones, M. R.; Rouillard, A. D.; Fernandez, N. F.; Duan, Q.; Wang, Z.; Koplev, S.; Jenkins, S. L.; Jagodnik, K. M.; Lachmann, A.; McDermott, M. G.; Monteiro, C. D.; Gundersen, G. W.; Ma’ayan, A. Enrichr: A Comprehensive Gene Set Enrichment Analysis Web Server 2016 Update. Nucleic Acids Research 2016, 44 (W1), W90–W97. https://doi.org/10.1093/nar/gkw377.

(36) Carlson, M. GO.Db: A Set of Annotation Maps Describing the Entire Gene Ontology. R Package Version 3.8.2. Bioconductor 3.10. 2019.

(37) Köhler, S.; Bauer, S.; Horn, D.; Robinson, P. N. Walking the Interactome for Prioritization of Candidate Disease Genes. The American Journal of Human Genetics 2008, 82 (4), 949– 958. https://doi.org/10.1016/J.AJHG.2008.02.013.

(38) Riley, N. M.; Coon, J. J. Phosphoproteomics in the Age of Rapid and Deep Proteome Profiling. Analytical Chemistry 2016, 88 (1), 74–94. https://doi.org/10.1021/acs.anal-chem.5b04123.

(39) Blatti, C.; Sinha, S. Characterizing Gene Sets Using Discriminative Random Walks with Restart on Heterogeneous Biological Networks. Bioinformatics 2016, 32 (14), 2167–2175. https://doi.org/10.1093/bioinformatics/btw151.

(40) Shieh, G. S. A Weighted Kendall’s Tau Statistic. Statistics & Probability Letters 1998, 39 (1), 17–24. https://doi.org/10.1016/S0167-7152(98)00006-6.

(41) Fey, D.; Croucher, D. R.; Kolch, W.; Kholodenko, B. N. Crosstalk and Signaling Switches in Mitogen-Activated Protein Kinase Cascades. Frontiers in Physiology 2012, 3. https://doi.org/10.3389/fphys.2012.00355.

(42) Arkun, Y.; Yasemi, M. Dynamics and Control of the ERK Signaling Pathway: Sensitivity, Bistability, and Oscillations. PLOS ONE 2018, 13 (4), e0195513. https://doi.org/10.1371/journal.pone.0195513.

(43) H. B. Mann; D. R. Whitney. On a Test of Whether One of Two Random Variables Is Stochastically Larger than the Other. The Annals of Mathematical Statistics 1947, 18 (1), 50– 60. https://doi.org/10.1214/aoms/1177730491.

(44) Benjamini, Y.; Hochberg, Y. Controlling the False Discovery Rate: A Practical and Powerful Approach to Multiple Testing. Journal of the Royal Statistical Society: Series B (Methodological) 1995, 57 (1), 289–300. https://doi.org/10.1111/j.2517-6161.1995.tb02031.x.

(45) Dow, L. E.; Elsum, I. A.; King, C. L.; Kinross, K. M.; Richardson, H. E.; Humbert, P. O. Loss of Human Scribble Cooperates with H-Ras to Promote Cell Invasion through Deregulation of MAPK Signalling. Oncogene 2008, 27 (46), 5988–6001. https://doi.org/10.1038/onc.2008.219.

(46) Nola, S.; Sebbagh, M.; Marchetto, S.; Osmani, N.; Nourry, C.; Audebert, S.; Navarro, C.; Rachel, R.; Montcouquiol, M.; Sans, N.; Etienne-Manneville, S.; Borg, J. P.; Santoni, M. J. Scrib Regulates PAK Activity during the Cell Migration Process. Human Molecular Genetics 2008, 17 (22), 3552–3565. https://doi.org/10.1093/hmg/ddn248.

(47) Pennock, S.; Wang, Z. A Tale of Two Cbls: Interplay of c-Cbl and Cbl-b in Epidermal Growth Factor Receptor Downregulation. Molecular and Cellular Biology 2008, 28 (9), 3020–3037. https://doi.org/10.1128/mcb.01809-07.

(48) Roepstorff, K.; Grandal, M. V.; Henriksen, L.; Knudsen, S. L. J.; Lerdrup, M.; Grøvdal, L.; Willumsen, B. M.; Van Deurs, B. Differential Effects of EGFR Ligands on Endocytic Sorting of the Receptor. Traffic 2009, 10 (8), 1115–1127. https://doi.org/10.1111/j.1600-0854.2009.00943.x.

(49) Neukamm, S. S.; Toth, R.; Morrice, N.; Campbell, D. G.; MacKintosh, C.; Lehmann, R.; Haering, H. U.; Schleicher, E. D.; Weigert, C. Identification of the Amino Acids 300-600 of IRS-2 as 14-3-3 Binding Region with the Importance of IGF-1/Insulin-Regulated Phosphorylation of Ser-573. PLoS ONE 2012, 7 (8). https://doi.org/10.1371/journal.pone.0043296.

(50) Gingras, A.-C.; Abe, K. T.; Raught, B. Getting to Know the Neighborhood: Using Proximity-Dependent Biotinylation to Characterize Protein Complexes and Map Organelles. Current Opinion in Chemical Biology 2019, 48, 44–54. https://doi.org/10.1016/j.cbpa.2018.10.017.

(51) Orre, L. M.; Vesterlund, M.; Pan, Y.; Arslan, T.; Zhu, Y.; Fernandez Woodbridge, A.; Frings, O.; Fredlund, E.; Lehtiö, J. SubCellBarCode: Proteome-Wide Mapping of Protein Localization and Relocalization. Molecular Cell 2019, 73 (1), 166–182.e7. https://doi.org/10.1016/j.molcel.2018.11.035.

(52) Maertens, A.; Tran, V. P.; Maertens, M.; Kleensang, A.; Luechtefeld, T. H.; Hartung, T.; Paller, C. J. Functionally Enigmatic Genes in Cancer: Using TCGA Data to Map the Limitations of Annotations. Scientific reports 2020, 10 (1), 4106. https://doi.org/10.1038/s41598-020-60456-x.

(53) Haynes, W. A.; Tomczak, A.; Khatri, P. Gene Annotation Bias Impedes Biomedical Research. Scientific Reports 2018, 8 (1), 1362. https://doi.org/10.1038/s41598-018-19333-x.

(54) Naro, C.; Sette, C. Phosphorylation-Mediated Regulation of Alternative Splicing in Cancer. International Journal of Cell Biology. 2013. https://doi.org/10.1155/2013/151839.

(55) Mukherji, M.; Brill, L. M.; Ficarro, S. B.; Hampton, G. M.; Schultz, P. G. A Phosphoproteomic Analysis of the ErbB2 Receptor Tyrosine Kinase Signaling Pathways. Biochemistry 2006, 45 (51), 15529–15540. https://doi.org/10.1021/bi060971c.

(56) Gaudet, P.; Dessimoz, C. Gene Ontology: Pitfalls, Biases, and Remedies. In The Gene Ontology Handbook; Dessimoz, C., Škunca, N., Eds.; Springer New York: New York, NY, 2017; pp 189–205. https://doi.org/10.1007/978-1-4939-3743-1_14.

