## Supplementary material for "Using Multilayer Heterogeneous Networks to Infer Functions of Phosphorylated Sites": SI

Figure S1. Optimization of RWHN parameters using the validation dataset.

Figure S2. Validation using randomly permuted networks.

Figure S3. Clustering of Francavilla et al. data.

Figure S4. Clustering of Ruprecht et al. data.

Figure S5. Clustering of Santra et al. data.

Table S1. PSP ON_PROCESS annotations, that describe biological function of sites, mapped to most closely related GOBP term.

Table S2. Summary of the experimental methods and results from each of the Case Study datasets.

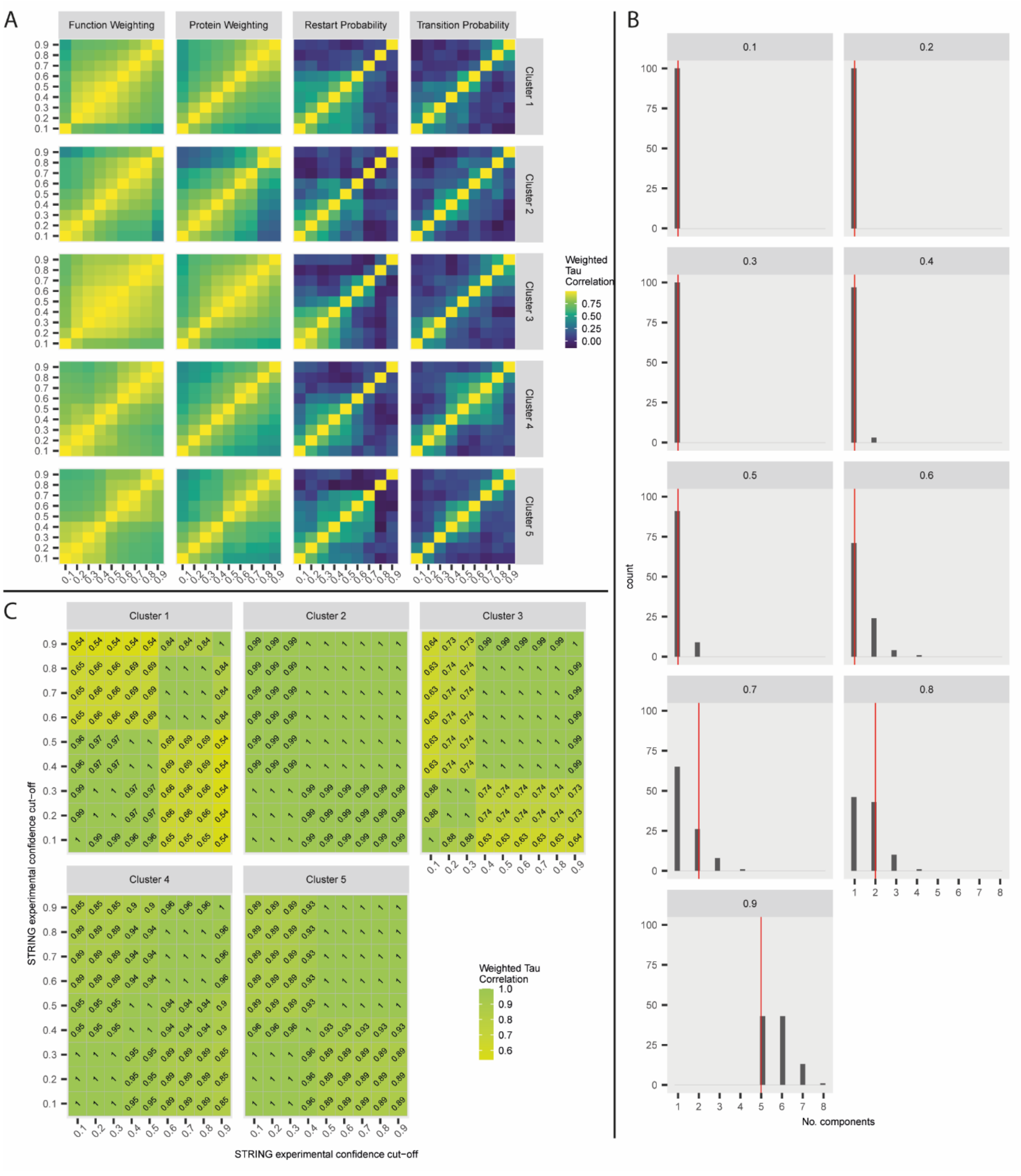


**Figure S1.** Optimization of RWHN parameters using the validation dataset. A) The parameters in the iterative RWHN calculation (Equation 8) – transition probability ($\lambda$), restart probability ($r$) – and the tunable weighting applied to the protein ($\eta_P$) or function ($\eta_F$) nodes of the multilayer heterogenous network were tested at values within the range of 0.1 – 0.9, changing one parameter whilst setting all others to 0.5. This was repeated with seed nodes set to all five clusters found in the validation dataset. We then calculated how frequently each GO term would be ranked at the same position when the value of each parameter was altered, indicating stability over the range of potential values. Each cell in the heatmaps indicates a GO Term at a rank, with the color indicating the percentage of times that term was given that rank over the range of parameters. B) Connectivity of randomly permuted networks based on the STRING PPI that forms the protein layer, with different confidence thresholds. As the confidence threshold increases above 0.7, the likelihood of the PPI being disconnected increased. The red line indicates the “true” PPI based on the validation data. C) Weighted Tau correlation of RWHN results with STRING PPI constructed with a range of confidence thresholds.

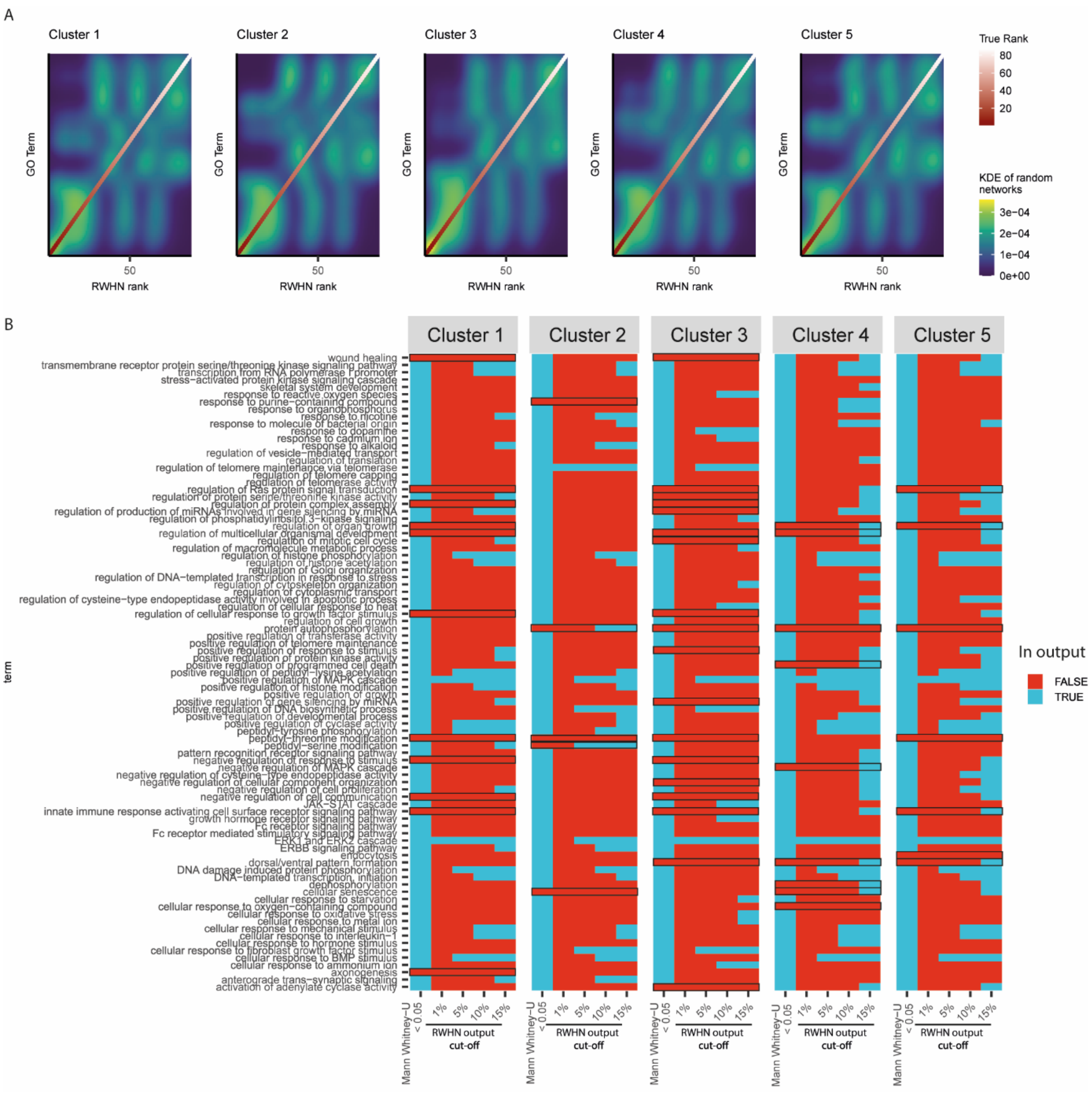


**Figure S2.** The validation network was permuted 100 times and RWHN run over each permutation with seed nodes set to sites found in clusters 1 to 5. A) KDE was calculated to estimate how frequently each term appeared at each rank in the random networks. The y-axis are reordered based on the "true" ranking of the GO terms (i.e., the ranking in the non-permuted network) for that cluster. If the random networks generated the same result as the actual network, we would expect the KDE plots to closely resemble the trend of the true rank. B) The probability of each term in the "true" result (i.e. RWHN results on the non-randomly permuted data) being ranked in that position for each cluster compared to the rankings from the randomly permuted networks was calculated using the Mann-Whitney U test ($p < 0.05$, with Benjamini & Hochberg correction). This was then compared to the RWHN output with different thresholds applied – the top 1%, 5%, 10% or 15% of terms. Those terms whose rankings did not differ from random are highlighted with a black outline.

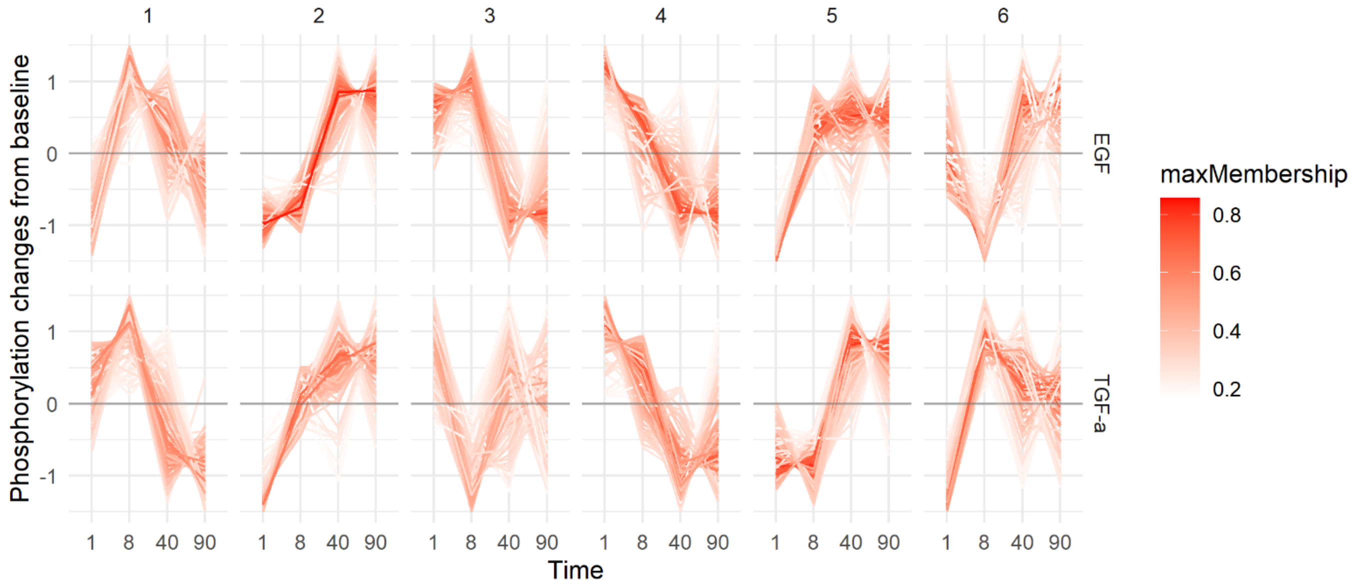


**Figure S3.** Phosphorylation levels over time of phosphorylated sites within each of the 6 clusters from the Francavilla et al. dataset that were regulated by EGF or TGF-α stimulation.

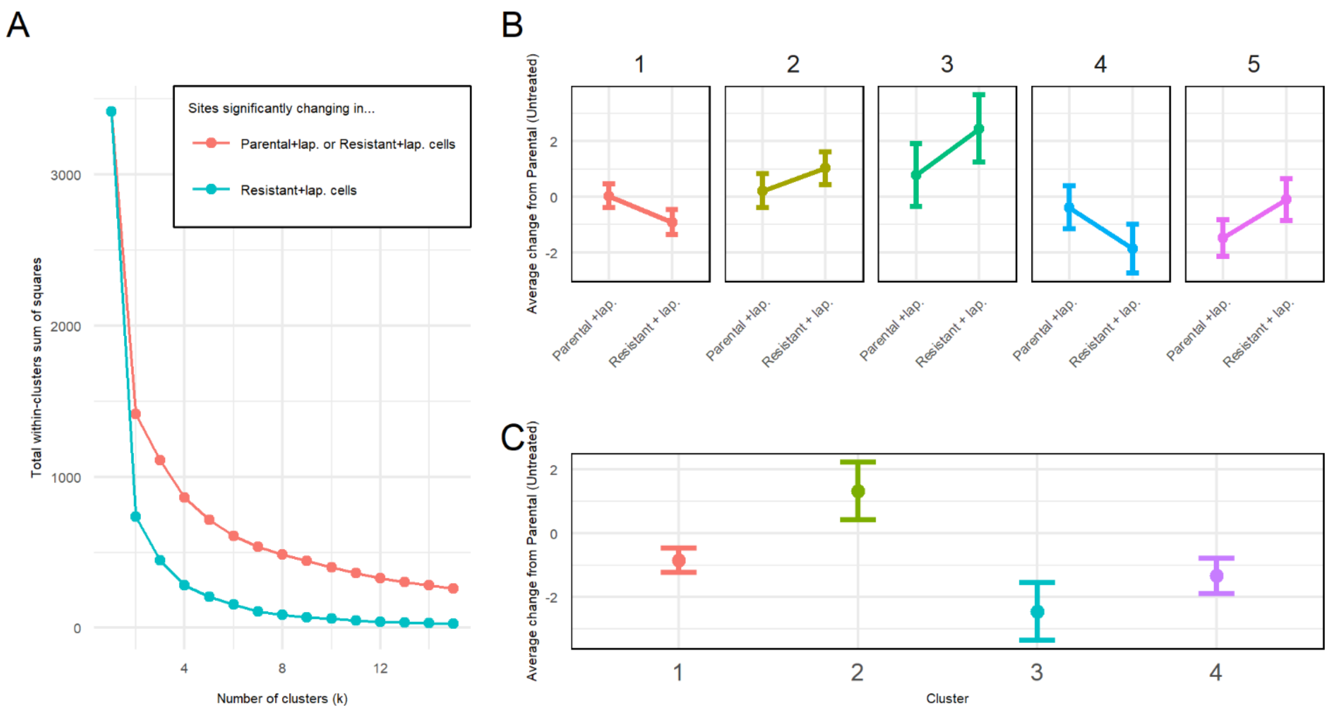


**Figure S4.** Clustering of regulated phosphorylated sites in the Ruprecht et al. dataset using k-means clustering. A) Elbow plot was used to determine the number of clusters (k) to group data into. The pink line represents sites regulated in lapatinib treated parental cells or resistant cells, whilst the blue line represents sites regulated in lapatinib-treated resistant cells only. B) Clustering of all lapatinib regulated sites in lapatinib-resistant and parental cell lines. C) Clustering of significantly changed sites in lapatinib-resistant cells.

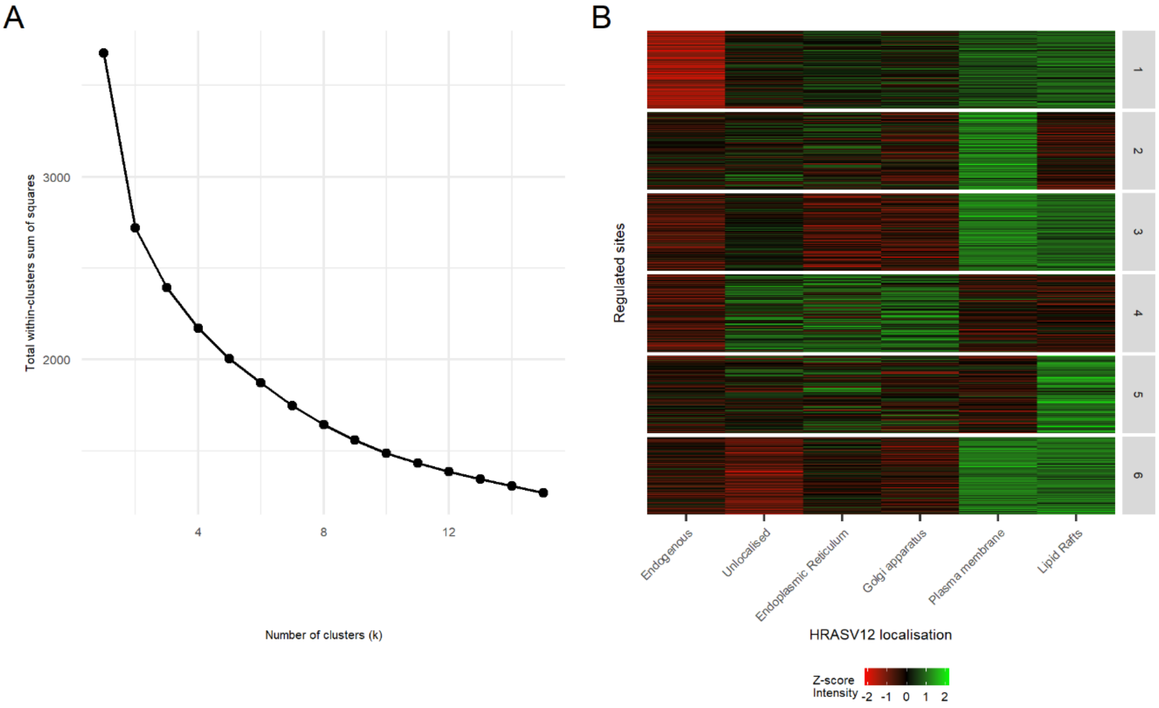


**Figure S5.** Clustering of regulated phosphorylated sites from the Santra et al. dataset using k-means clustering. A) Elbow plot used to determine the number of clusters (k) to group the data into. B) Clustering of regulated sites into the selected 6 clusters

**Table S1.** PSP ON_PROCESS annotations, that describe biological function of sites, mapped to most closely related GOBP term

| PSP "ON_PROCESS" annotation | Mapped GOBP term | GOID |
|---|---|---|
| apoptosis | apoptotic process | GO:0006915 |
| autophagy | autophagy | GO:0006914 |
| cell adhesion | cell adhesion | GO:0007155 |
| cell cycle regulation | cell cycle | GO:0007049 |
| cell differentiation | cell differentiation | GO:0030154 |
| cell growth | cell growth | GO:0016049 |
| cell motility | cell motility | GO:0048870 |
| chromatin organization | chromatin organization | GO:0006325 |
| cytoskeletal reorganization | cytoskeletal reorganization | GO:0007010 |
| DNA repair | DNA repair | GO:0006281 |
| endocytosis | endocytosis | GO:0006897 |
| exocytosis | exocytosis | GO:0006887 |
| RNA splicing | RNA splicing | GO:0008380 |
| RNA stability | regulation of RNA stability | GO:0043487 |
| signaling pathway regulation | signal transduction | GO:0007165 |
| transcription | Gene expression | GO:0010467 |
| translation | translation | GO:0006412 |

**Table S2.** Summary of the experimental methods and results from each of the Case Study datasets.

| Case Study | Public-ation | MS Quantification method / MS Instrument | Sample type | Experimental Conditions | Brief description of main findings |
|---|---|---|---|---|---|
| 1 | Francavilla et al. 2016 20 | SILAC / Q-Exactive | Cell culture: HeLa, serum starved | Unstimulated HeLa; Stimulation with EGF or TGF-α 100 ng/mL for 1, 8, 40 and 90 minutes, performed in duplicate | EGFR degradation upon stimulation with EGF was accompanied by phosphorylation of the late-endosome/lysosome (ie degradative pathway) marker Rab7 within 40 minutes of stimulation, and transient MAPK/ERK signalling. TGF-α stimulation, meanwhile, induced EGFR recycling to the plasma membrane through the recruitment of RCP prior to internalisation. This occurs alongside sustained MAPK/ERK signalling and a greater mitogenic and migratory response. |
| 2 | Ruprecht et al. 2017 21 | Dimethyl labelling / Q-Exactive Plus | Cell culture: Parental BT-474 and | Untreated parental BT-474; parental | Lapatinib treatment in parental BT-474 strongly altered splicing processes |

| | | | | | |
|---|---|---|---|---|---|
| | | | lapatinib resistant clone BT-474-J4 | BT-474 treated with 1 µmol/L lapatinib for 30 minutes; lapatinib resistant clone BT-474-J4 continuously cultured in 1 µmol/L lapatinib | and glycolytic processes via phosphorylation. Metabolism was recovered in resistant cells through phosphorylation-mediated reprogramming (e.g. phosphorylation of LDHA, ENO1, ALGOA, PFKP and GAPDH). Recovery of the transcriptional programme was not investigated. AXL/PI3K signalling had been previously reported to be downregulated by lapatinib treatment but recovered in resistant cells, and that was reconfirmed here. |
| 3 | Santra et al. 2019 22 | Label-free / LTQ-Orbitrap Elite | Cell culture. HeLa cells expressing endogenous HRAS, flag-HRAS, or flag-HRASV12 | Flag-IP to isolate interactors of the different protein baits. | Whilst protein interaction data showed HRASV12 has the highest number of interactors at the ER and DM, most of its effects on canonical downstream pathways such as MAPK and PI3K were initiated |

| | | | | | |
|---|---|---|---|---|---|
| | | | constructs targeted to one of the following subcellular organelles: disordered plasma membrane (DM), plasma membrane lipid rafts (LR), Golgi apparatus (GA) or endoplasmic reticulum (ER). | | from the DM and LR. It was found that the RAC and PAK pathways, along with HRAS' influence on the cell cycle, were mediated from the ER. A potent effect of HRASV12 localisation was demonstrated by the expression of TP53, which was overexpressed when HRASV12 was localised to the GA or DM and under expressed when it was localised to the ER. This had a knock-on effect on TP53's transcriptional targets. Finally, the localisation of HRASV12 to the ER was more strongly associated with migration than the other compartments, and predicted to be mediated via ERK signalling. |
